# Lords of the flies: Dipteran migrants are diverse, abundant and ecologically important

**DOI:** 10.1101/2024.03.04.583324

**Authors:** Will L. Hawkes, Myles H.M. Menz, Karl R. Wotton

**Affiliations:** Centre for Ecology and Conservation, University of Exeter, Cornwall Campus, Penryn, United Kingdom; College of Science and Engineering, James Cook University, Townsville, QLD 4811, Australia; Swiss Ornithological Institute, Sempach, 6204, Switzerland

## Abstract

Insect migrants are hugely abundant and recent studies have identified Diptera as the major component of many migratory assemblages, often totalling up to 90% of all individuals. Despite this, studies into their migratory behaviour have been widely eschewed in favour of the more ‘charismatic’ migrant insects such as butterflies, dragonflies, and moths. Here we review the available literature on Dipteran migration and identify 13 lines of evidence that we use to determine migratory behaviour. Using this approach, we find species from 60 out of 130 Dipteran families that show evidence of migration, with Syrphidae fulfilling 12 of these criteria, followed by the Tephritidae with 10. In contrast to these groups, 22 families fulfilled just two lines of evidence or fewer, underlining the need for more research into the migratory characteristics of these groups. In total, 622 species of Diptera were found to have migratory behaviour (0.5% of the total Dipteran species count), a figure rising to 3% for the Syrphidae, a percentage mirrored by other animal taxa such as butterflies, noctuid moths, and bats. Research was biased to locations in Europe (49% of publications) and while vast regions remain understudied, our review identified major flyways used by Dipteran migrants across all biogeographic realms. Finally, we detail the ecological and economic roles of these migrants and review how these services are being affected by anthropogenic change through population declines and phenological shifts. Overall, this review highlights how little is known about Dipteran migration and how vital their migratory behaviour may be to the health of global ecosystems.

## Introduction

Each year, huge numbers of insects migrate globally to exploit seasonally available resources to increase their reproductive output, and/or escape habitat deterioration, e.g., due to temperature change, disease risk, food quality, or to seek overwintering sites (Chapman et al., 2015; Dingle, 2014; Satterfield et al., 2020). Some insects are known to migrate hundreds and even thousands of kilometres in a single journey (Hobson et al., 2012), utilising the sun as a compass and favourable winds to power their journeys (Gao et al., 2020; Knoblauch et al., 2021; Massy et al., 2021; Menz et al., 2022; Stefanescu et al., 2013). Studies of insect migration have mainly focussed on the larger, more charismatic insects (Menz et al., 2022; Stefanescu et al., 2013; Wikelski et al., 2006) or agriculturally important species (Jia et al., 2022; Jones et al., 2019; Li et al., 2020). Few have systematically analysed whole migratory assemblages. However, the studies that do exist have revealed a major group of migrants that remain hugely understudied and that are of great ecological importance: the Diptera (Hawkes et al., 2022, 2024).

The Diptera are a huge Order of insects, consisting of over 125,000 described species, although over 1 million species are estimated to exist (Wiegmann et al., 2011). Dipteran migration behaviour is poorly known and little studied, despite mass occurrences being frequently observed, including potentially two of the ten Plagues of Egypt described in the book of Exodus: gnats and dog-flies (Brenton, 1844). Likewise, in Serbian mythology, a legend concerning the death of a she-demon called an Ala notes the spring arrival of a plague of Golubatz (*Simulum colombaschense*) flies from the rotting corpse (Караџић, 2005). This legend too suggests its truth lies within insect migration (Babic et al., 1935). Recent systematic studies of insects passing through migratory hotspots have shown that the Diptera often comprise nearly 90% of the individuals found in migratory assemblages in certain locations (Hawkes et al., 2022, 2024). Ecological assessments of these species suggest that these flies play a huge range of ecological roles of importance to both the anthropogenic and natural world (Doyle et al., 2020; Hawkes et al., 2022; Wiegmann et al., 2011). However, when compared to the migration of vertebrates and some other insect groups (e.g., the Lepidoptera) very little is known and what information there is, is highly dispersed (Chowdhury et al., 2021; Dingle, 2014).

In this review we collate all the known information about dipteran migration globally including which Families and species display migratory behaviour. We use this information to identify potential flyways, describe the ecological roles of these migrants, and explore the impacts that anthropogenically induced climate change may have on their migration.

## Defining migration

A widely used definition of migration is one based on behavioural characteristics: ‘Migratory behaviour is persistent and straightened-out movement effected by the animal’s own locomotory exertions or by its active embarkation on a vehicle’ (Kennedy, 1985). It depends on some temporary inhibition of station keeping responses but promotes their eventual disinhibition and recurrence’ (Kennedy, 1985). Dipteran migrants, and migratory insects in general, are subject to various viewpoints as to what constitutes migration (e.g., butterfly migration, Chowdhury et al., 2021). Therefore, similarly to the recent butterfly migration review (Chowdhury et al., 2021), we use the broad behavioural definition of migration quoted above, while recognising that we can only be certain of migratory behaviour from a few species. Instead of this representing a failure of the definition, we believe it is a result of a lack of research into the migratory behaviour of Diptera. This broader viewpoint utilised in this review is hoped to establish an initial baseline for future research into the migratory behaviour of Diptera.

## Literature Search

Google Scholar, Web of Science and PubMed were searched to determine which of the Dipteran families show migratory behaviour based on at least one line of evidence which suggests some level of migratory behaviour: Seasonal back and forth movement, long distance flight, seasonally appropriate directed movement, inability to develop in trapped habitat, ability to choose favourable winds, mass arrival, capable of high-altitude flight, populations with a high rate of gene flow, strong flight capabilities (tethered flight mill), orientation within a flight simulator, Physiological/morphological changes in the migratory phenotype, seasonal appearance of a disease, unable to overwinter (in any state) in location (see Table 1).

**Table 1.** Migratory criteria. Criteria 1-4 form the ‘core 4’ most often reported migratory characteristics.

| Migratory criteria | Description | Example references |
| --- | --- | --- |
| (1) Seasonal back and forth movement | Perhaps the strongest indicator of migration, the insects are observed during the springtime and then again in the autumn season. This can be evidenced by peaks in numbers in different migratory seasons (through radar data/citizen science recording etc.) or actively seeing the insects moving purposefully in one direction during | Florio et al., 2020 |
|  | one season, and then back the opposite direction later in the year. |  |
| (2) Long-distance flight | Long-distance flight is important for migratory insects to escape unfavourable habitats. | Hawkes et al., 2022 |
| (3) Seasonally appropriate directed movement | Directed movement of an insect in a seasonally appropriate direction (e.g. higher latitudes in spring, lower latitudes in autumn) suggests a preferred flight detection. | Lack & Lack, 1951 |
| (4) Inability to develop in trapped habitat | Larvae are incapable of developing due to unfavourable seasonal climate. This suggests that the adult insects must move away from their current location to lay their eggs in order for their young to survive. | Ashmole et al., 1983 |
| (5) Ability to choose favourable winds | An important factor in insect migration as the winds are used to power their migrations. | Gao et al., 2020 |
| (6) Mass arrival | Migratory flies often arrive in large numbers at the same time. | Hawkes et al., 2024 |
| (7) Capable of high-altitude flight | To migrate, flies will take advantage of higher altitude wind currents. Additionally, there is little reason for insects to be found consistently at altitude if they are not attempting to move larger distances. | Chapman et al., 2004 |
| (8) Populations with a high rate of gene flow | This suggests a high level of movement between populations by individuals. | Mignotte et al., 2021 |
| (9) Strong flight capabilities (tethered flight mill) | To migrate long-distances, insects must have strong flight capabilities. This can be evidence by their performance in a flight mill. | Nilssen & Anderson, 1995 |
| (10) Orientation within a flight simulator | A preferred, seasonally advantageous flight direction in a flight simulator is indicative of migratory behaviour. | Massy et al., 2021 |
| (11) Physiological/morphological changes in the migratory phenotype | This includes any physiological or morphological changes associated with a migratory phenotype. Including delaying the development of reproductive organs, or changes in morphology between resident and migratory generations. | Doyle et al., 2023 |
| (12) Seasonal appearance of a disease | If the insects are associated with a seasonal appearance of a disease, then it is likely that they are acting as vectors - bringing the disease from faraway locations. | Nabeshima et al., 2009 |
| (13) Unable to overwinter (in any state) in location | The adult insects were trapped in a region where they are not capable of surviving the winter (e.g., at high latitudes, above oceans, in high mountain passes etc.). | Ashmole et al., 1983 |

To obtain an initial overview, performed up to March 2024, the search results were filtered to include the words ‘Diptera’ and ‘Migration’ anywhere in the manuscript, without ‘larvae’ and ‘cell’ and ‘development’ to exclude evolutionary development studies. ‘Dispersal’ was also avoided as this swamped the literature with papers documenting small scale movements of Diptera (∼<300m). This methodology yielded 6200 results and the first 1000 papers were carefully analysed for relevancy. A provisional list of migratory Dipteran families (‘X’) was obtained from these papers before Google Scholar was then used to search for specific information on each of these families using the term ‘X migration’. In Web of Science, the search term ‘Diptera Migration’ without the term ‘cell’ yielded 700 results. In PubMed the same search criteria returned 993 results. A specific search of Dipteran families was also carried out for both Web of Science and PubMed databases. To collect results that may not be included in online search databases due to age, further searches were performed within books such as ‘Mechanisms of Insect Dispersal: Migration and Dispersal of Insects by Flight’ (Johnson 1969), ‘Insect Migration’ (Williams, 1958) and in the reference lists of relevant articles. Ultimately, suggestions that saturation was close to being reached occurred when repeated and irrelevant works were found during literature searches of Google Scholar, Web of Science, and PubMed. Searches were conducted up to October 2022 and in total 193 relevant articles were identified.

## Prevalence of migration

In total, we found that ∼47% of all Dipteran families (60/130) had evidence of migratory behaviour from at least one species. A detailed table of evidence including the papers used can be found in the supplementary file Table S1. Of the 193 papers that contained evidence of Dipteran migration, 93 (or 48%) provided evidence of Syrphidae (hoverflies) migration, making them the most well-studied of the migratory Dipteran families. The Syrphidae also fulfilled the most migratory criteria of any Family: 12/13 criteria (Table 1, Figure 1, and Table S1) missing only the ‘seasonal appearance of a disease’ criteria. The Culicidae (mosquitoes) were the second most studied with 32 (or 17%) of the papers and fulfilled the third most migratory criteria behind the Tephritidae (fruit flies) with 10/13, and alongside Muscidae (house flies), and Calliphoridae (blow flies and screw worms) and Chloropidae (grass flies): 9/13 (Figure 1). Chloropidae are miniscule creatures (∼2mm in length) yet have been recorded showing a core set referred to here as the ‘core four’ of ‘Seasonal back and forth movements’, ‘long-distance flight’, ‘seasonally adaptive directed movements’ and ‘inability to develop in trapped habitat’ suggesting strong migratory behaviour. Additionally, a study in a high-altitude Pyrenean pass showed their ability to choose favourable winds (Hawkes et al., 2024), while a North American aerial study found individuals flying at over 1,500m in elevation (Glick, 1939).

**Figure 1.**
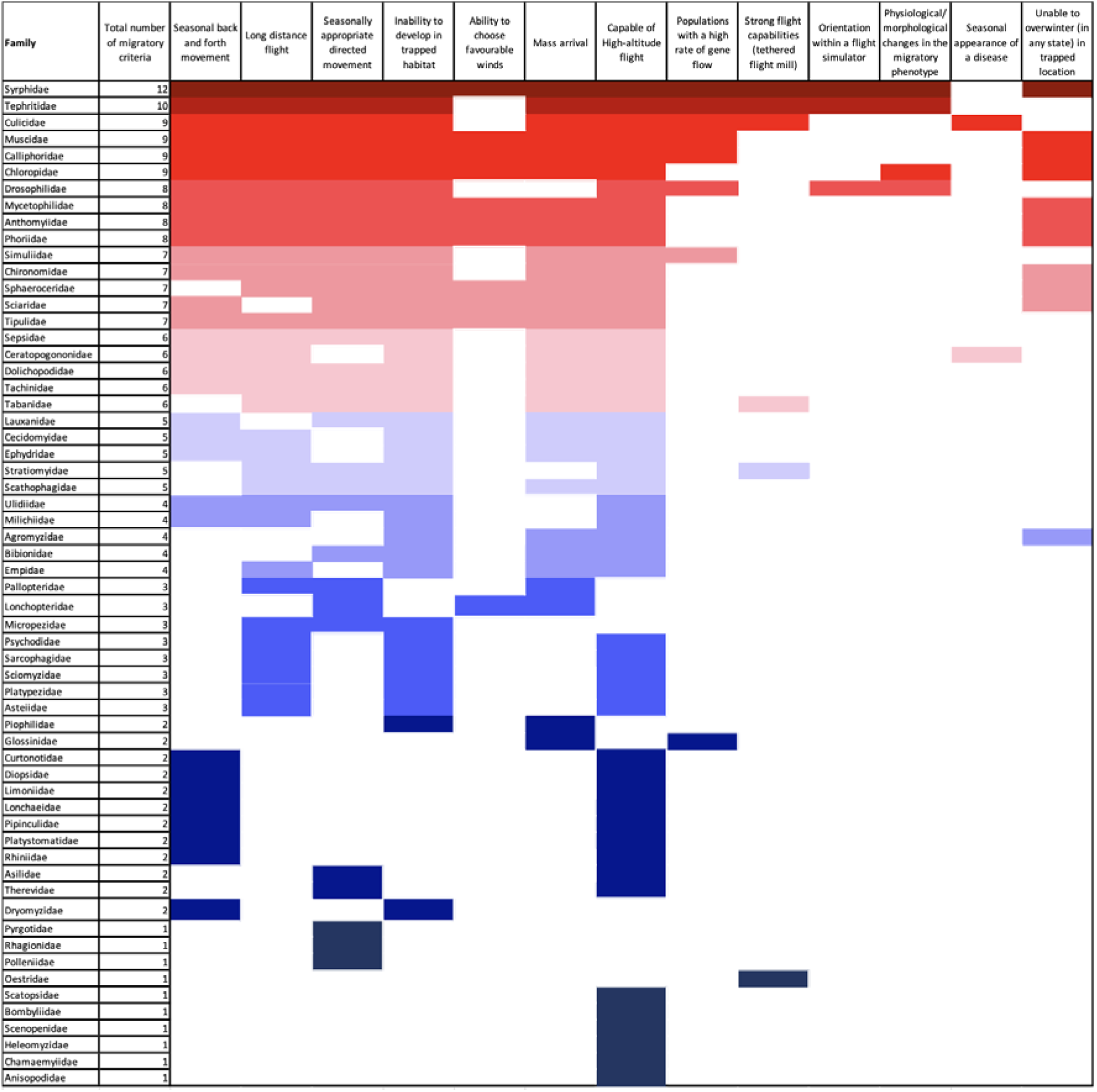
The known migratory criteria fulfilled by the 60 identified migratory families of Diptera. Heat map colours indicate the number of migratory criteria identified for each family with red being the most (12 criteria fulfilled) and dark blue the least (one criteria fulfilled).

Drosophilidae (fruit flies), Mycetophilidae (fungus gnats), Anthomyiidae (root maggots), Phoriidae (scuttle flies) fulfilled 8/13 migratory criteria, all fulfilling the ‘core four’. *Delia platura* (Anthomyiidae) have been recorded in their millions numbers migrating from the Middle East to Cyprus along a northeast trajectory during the springtime, a journey representing at least 105m of ocean crossing (Hawkes et al., 2022).

Simuliidae (black flies), Chironomidae (non-biting midges), Sphaeroceridae (small dung flies), Sciaridae (black fungus gnats), Tipulidae (crane flies) fulfilled 7/13 migratory criteria. Simuliidae, Chironomidae, and Tipulidae fulfilled the ‘core four’ criteria, while Sphaeroceridae and Sciaridae missed ‘seasonal back and forth movement’ and ‘long-distance movement’ respectively. Tipulidae, for example, showed seasonal back and forth movement at high altitude above Mali, as well as evidence of long-distance flight after being found on oil rigs in the North Sea (Hardy & Cheng, 1986), or trapped in nets from ships in the Gulf of Mexico (Keaster et al., 1996). Additionally, Gatter (1977), recorded Tipulidae utilising favourable winds in large numbers migrating through the mountains of southwest Germany.

Five of the 60 families fulfilled 6/13 migratory criteria; Sepsidae (ant-like scavenger flies), Ceratopogonidae (biting midges), Dolichopodidae (long-legged flies), Tachinidae (tachinid flies), and Tabanidae (horse flies). All families bar Ceratopogonidae (missing seasonally adaptive movement) and Tabanidae (missing seasonal back and forth movement) fulfilled the ‘core four’ criteria. Ceratopogonidae flies, like many others, were recorded showcasing seasonal back and forth movement at high altitude above Mali (Florio et al., 2020). They have been shown to be capable of long-distance flight by being recorded in the middle of the Gulf of Mexico where of course none of their larvae could survive (Keaster et al., 1996). Fascinatingly this family has also been shown to be a vector of livestock diseases such as bluetongue and schmallengberg viruses (Mignotte et al., 2021).

Six of the 60 families fulfilled 5/13 migratory criteria. These were the Lauxanidae (Lauxaniid flies), Cecidomyidae (gall midges), Ulidiidae (picture-winged flies), Ephydridae (shore flies), Stratiomyidae (soldierflies), and Scathophagidae (dung flies). Of these six, only Ulidiidae fulfilled all the ‘core four’ criteria. They were recorded showing seasonal back and forth movement above Mali (Florio et al., 2020), been trapped at sea in the Gulf of Mexico showing long-distance flight in an area where their larvae cannot develop, and seasonally adaptive directed movement through the Pass of Portachuelo in Venezuela (Beebe, 1951).

A further four families filled 4/13 migratory criteria: Milichiidae (jackal flies), Agromyzidae (leaf-miner flies), Bibionidae (march flies), and Empidae (dance flies). Again, none recorded the ‘core four’ criteria yet all recorded being trapped in areas where their larvae could not develop. Bibionidae, for example, were recorded after a migration fallout in the snowfields of the Cairngorms (Ashmole et al., 1983). Additionally, Bibionidae have been recorded moving purposefully through the Pass of Portachuelo, Venezuela, in large numbers. On May 29^th^, 1948, Beebe (1951) noted a *Bibio* sp. moving through the pass accompanied by a ‘veritable mist of others’.

Pallopteridae (flutter-winged flies), Lonchopteridae (spear winged flies), Micropezidae (stilt-legged flies), Psychodidae (owl midges), Sarcophagidae (flesh flies), Sciomyzidae (snail killing flies), Platypezidae (flat-footed flies), and Asteiidae (asteiid flies), were the eight families which recorded 3/14 migratory criteria. ‘Long distance flight’, ‘incapable of developing in trapped location’, and ‘capable of high-altitude flight’ were the commonest criteria met with many of the families found in the middle of the Gulf of Mexico (Sparks et al., 1986; Wolf et al., 1986) and at high altitude above North America by Glick (1939).

The largest group contained 12 of the 60 families and fulfilled 2/13 migratory criteria. These families were Curtonotidae (small dung flies), Diopsidae (stalk-eyed flies), Limoniidae (crane flies), Lonchaeidae (lance flies), Pipinculidae (big-headed flies), Platystomatidae (signal flies), Rhiniidae (Rhiniid flies), Asilidae (robber flies), Therevidae (stiletto flies), and Dryomyzidae (Dryomyzid flies). ‘High-altitude’ flight was the most common criteria met. Asilidae were recorded at medium-high altitude above North America (Glick, 1939) and also showed seasonally appropriate directed movement through the Portachuelo Pass in Venezuela (Beebe, 1951).

Finally, 11 of the 60 families fulfilled just 1/13 migratory criteria. These families were, Pyrgotidae (picture-winged flies), Rhagionidae (snipe flies), Pollenidae (cluster flies), Oestridae (warble flies), Scatopsidae (dung midges), Bombylidae (beeflies), Scenopenidae (window flies), Heleomyzidae (spiny-winged flies), Chamaemyiidae (Chamaemyid flies), and Anisopodidae (wood gnats). ‘High-altitude flight’ was the commonest criteria met, with Bombyliidae recorded at 60m in the air (Glick, 1939). Finally, Oestridae (bot and warble flies), also only met one of the criteria: ‘strong flight capabilities on a tethered flight mill’ with a singular paper showing that the reindeer warble fly *Hypoderma tarandi* can fly for 31.5 hours, with a longest continual flight of 12 hours (Nilssen & Anderson, 1995). This behaviour must play a role in the insect’s life history, likely for following their host species reindeer on their own great migrations, but no further supporting evidence currently exists.

Evidence for migratory was behaviour was found for 622 species (see Supplementary File S1 for a full species list), making around 0.5% of identified Dipteran species migratory. However, in the Syrphidae, 212 of the known 6000 species migrate, equal to 3.5% of the species in this family (see Figure 2). Interestingly, within the butterflies, a well-studied group of migratory insects, 3% of all species have been diagnosed as migratory (Chowdhury et al., 2021) and the same 3% is true for the noctuid moths (Alerstam 2011) and bat species (Fleming et al., 2003). While this may suggest an emerging pattern across taxa, more research is certainly needed.

**Figure 2.**
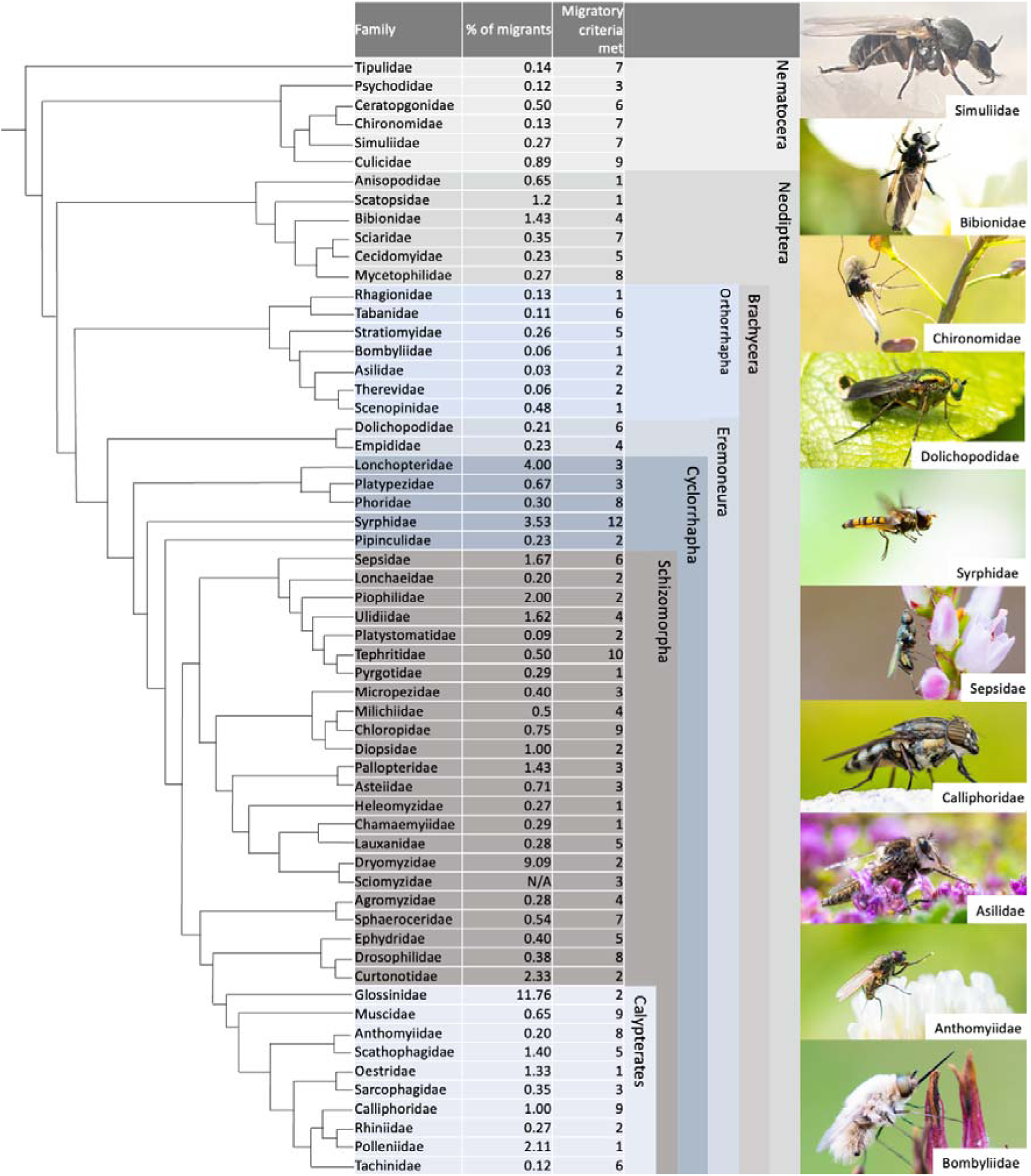
Migratory Diptera phylogeny and percentage of migratory species in each family based on Wiegmann et al., 2011. All photos ©Will Hawkes, apart from Simuliidae (©Mehmet Akif Suna).

## Mechanisms of fly migration

To migrate long distances, flies rely on a variety of mechanisms to both power and orientate themselves on their journeys. Studies performed at migratory hotspots focussing on the migratory criteria needed for fly migration to occur suggest that warmer temperatures, dry conditions, and the presence of winds favourable to the preferred migratory direction are important (Hawkes et al., 2022, 2024 under review). A radar study across southern Britain showed that Syrphidae actively select winds in the autumn which aid southward migration (Gao et al., 2020). During the springtime too, Syrphidae and other Diptera have been recorded arriving at locations in Europe on winds from the south (Gao et al., 2020; Hawkes et al., 2022; Hawkes et al., 2022). Flying higher in favourable tailwinds allows migratory insects to fly faster than their self-powered airspeed (Chapman et al., 2016; Gao et al., 2020). The speed of migratory Syrphidae above southern England have been recorded in the springtime at 11.2 m/s, and 9.8 m/s in the autumn (Gao et al., 2020), only a little slower than the speeds of nocturnal migrating moths (spring: 16.57 m/s, autumn: 13.75 m/s) and songbirds (spring: 13.48 m/s, autumn: 12.14 m/s) (Chapman et al., 2016).

Because the insects can select favourable winds, this points to the presence of a compass system within the migratory Diptera. In tethered flight simulator experiments, *Drosophila melanogaster* have been shown to maintain a constant flight heading utilising the sun and polarised light patterns (Warren et al., 2019; Weir & Dickinson, 2012). However, these headings are arbitrary with respect to a simulated sun and there is no evidence of time compensation as the sun moves across the heavens (Warren et al., 2019; Weir & Dickinson, 2012). A flight simulator experiment performed on two species of Syrphidae (*Scaeva pyrastri* and *S. selenitica*) caught while migrating through the Pyrenees during the autumn, showed that these larger flies do have a time compensated sun compass enabling them to maintain their preferred migratory heading even as the sun moves throughout the day (Massy et al., 2021). The status of such a compass in other migratory Diptera remains to be investigated.

In addition to using environmental cues, Diptera also undergo changes in their physiology during migration. These changes allow them to store energy and prepare for the long journey ahead. For example, flies will increase their fat stores before migrating, which provides them with the energy they need to fly long distances (Hondelmann & Poehling, 2007). A study into the genomes of non-migratory summer individuals and migratory autumn individuals trapped in a high-altitude Pyrenean pass, revealed over 1500 genes showing strong evidence for differential expression between the generations (Doyle et al., 2022). Analyses of these genes reveal a remarkable range of roles in metabolism, muscle structure and function, hormonal regulation, immunity, stress resistance, flight and feeding behaviour, longevity, reproductive diapause, and sensory perception, all of which are key traits associated with migration and migratory behaviour (Doyle et al., 2022).

## Global distribution and Flyways

We found a globally widespread distribution of migratory behaviour in Diptera (Figure 3). Records were recovered from all continents including, surprisingly Antarctica, where the Calliphorid *Calliphora croceipalpis,* was identified as likely migrant on the sub-Antarctic Marion Island, 1700km away from South Africa, the closest non-snow-covered landmass (Chown K, 1994). Our data points to a bias of European migration records, which make up 49% of publications (Figure 3), followed by Asia (13%), North America, Africa, and Australasia (all 10%). These distributions point to important flyways which we discuss in the next sections.

**Figure 3.**
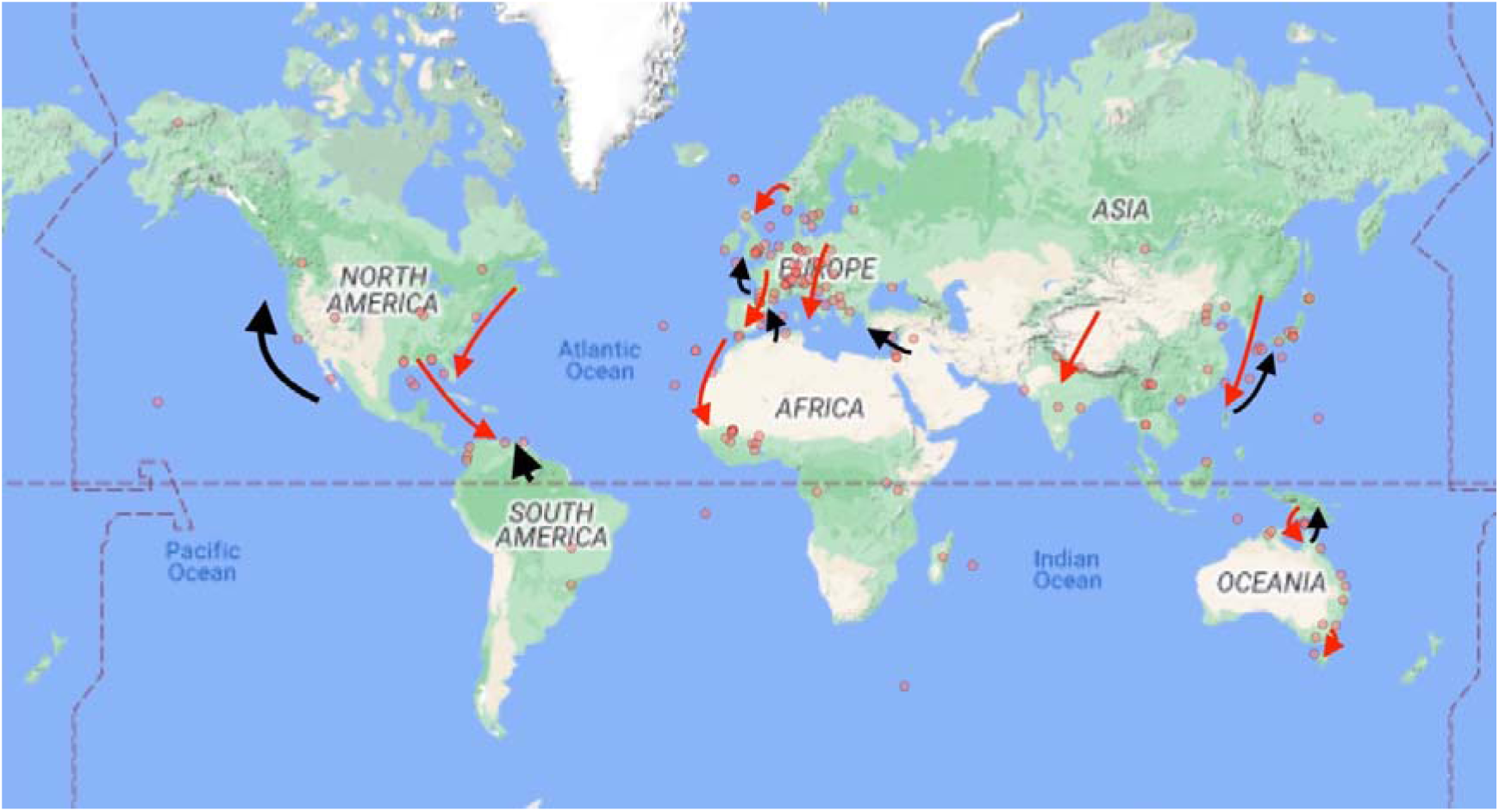
The geographic distributions of Dipteran migration studies and migratory flyways. Black arrows represent suggested northward migration routes, Red arrows represent southward routes. Red dots represent the locations of the Dipteran migration studies identified here.

### Eastern and Western seaboard flyways of North America

On the western seaboard of North America, southward migration of Diptera during the Autumn season was recorded multiple times between 1915 and 1926 (Shannon, 1926). Species listed were Calliphoridae*: Cochlyomia macellaria, Calliphora vicina, Phormia regina*; Muscidae: *Stomoxys calcitrans*, and Syrphidae: *Eristalis tenax,* moving south “in their thousands” (Shannon, 1926). No further observations being made in the 90+ years since, and the status of these movements is currently unknown (Menz, Brown, et al., 2019). However, recent isotopic studies on the Syrphid *Eupeodes americana* suggest that these flies are capable of moving up to 3000 km from Canada to Alabama, indicating that the flyway down the eastern part of North America is long distance and may still be well-utilised by migratory Diptera (Clem et al., 2023) along with other migratory insects (Howard & Davis, 2009; Wikelski et al., 2006). In contrast, only one movement has been identified on the western seaboard, a northward movement of presumed *Eupeodes* sp. Syrphidae numbering in the hundreds of thousands in just half an hour, recorded on the west coast of California in April 2017 (Menz, Brown, et al., 2019). Although no further observations exist, it is likely that migratory Diptera regularly move north in the springtime and southwards in the autumn to exploit seasonal resources in North America. Citizen science data for the *Eupeodes* genus suggests that these flies move from 35° N latitude during the winter months to 65° N during the springtime, suggesting seasonal long-distance movement along this flyway (Menz, Brown, et al., 2019).

### Cross Caribbean flyway

Many Nearctic bird species are known to migrate to the neotropical regions of South America to overwinter (Sainz-Borgo et al., 2020). Alongside many North American migratory birds, vast quantities of migratory insects have been recorded flying south in the Autumn through the Pass of Portachuelo, Venezuela (Beebe, 1951). This pass runs N/S and opens towards the Caribbean Sea, collecting any insects flying across the ocean. In the late 1940s, insect migration was so plentiful through the pass that the researchers had to wear glasses to protect their eyes from the abundant swarms (Beebe, 1951). In this pass, 17 families of Diptera were recorded, all moving North to South (Beebe, 1951). Although no insect related studies have occurred in the pass of Portachuelo since, more recent studies have recorded a variety of migratory Diptera (26 families) alighting on ships and oil rigs in the centre of the Gulf of Mexico, indicating that Diptera migration likely occurs across the entirety of the Caribbean Sea (Keaster et al., 1996; A. N. Sparks et al., 1986).

### Western European flyway

The Western European flyway is perhaps the best studied of all the flyways of migratory Diptera, although it is telling that the flyway is still hugely understudied. Long-term, whole assemblage, studies have been performed on migratory Diptera from this region from suction traps in the UK, to mountain hotspot studies in Germany, the French/Swiss Alps, the Pyrenees, and the Czech Republic (Aubert et al., 1976; Chapman et al., 2004; Gatter et al., 2020; Hlaváček et al., 2022; Lack & Lack, 1951; Snow & Ross, 1952; Williams et al., 1956, Hawkes et al. 2024), and there have been many observations of migratory Diptera made from locations in the far north such as Norway and within the North Sea, as well as south to the tip of Gibraltar (Ebejer & Bensusan, 2010; Hardy & Cheng, 1986; Jensen, 2001; Nielsen et al., 2010). A four-year study at a Pyrenean mountain pass in the autumn season revealed 12 families of migratory Diptera migrating south (Hawkes et al., 2024). Radar studies have revealed the directional movements of migratory Diptera, detailing a SSW bias in their autumnal movements (Chapman et al., 2010; Gao et al., 2020; Odermatt et al., 2017). This suggests that migratory Diptera found in Western Europe in the Autumn will be funnelled down into the Iberian Peninsula from large swathes of Europe, before potentially crossing into northern Africa via the straits of Gibraltar (Ebejer & Bensusan, 2010).

The majority of Dipteran migration studies have been performed in the autumn, but hints at their springtime routes are available. Large numbers of migratory Syrphidae have been found in the dunes during the springtime at Gibraltar, having just crossed the straits from Africa (Ebejer & Bensusan, 2010). In 2022, large numbers of migratory Diptera, primarily Syrphidae, were found washed up on a beach in SW France, wind analyses suggesting they were moving north over the Mediterranean before drowning due to a storm (Fisler & Marcacci, 2022). In the same year, large numbers of multiple species of Syrphidae were found to have arrived on the Isles of Scilly, UK, wind analysis suggesting that they took off over 200 km away in western France (Hawkes et al., 2022). *Culicoides obsoletus* (Ceratopogonidae) fly populations, which spread bluetongue and Schmallenberg viruses, were found to have high levels of gene flow and no genetic structuring at the scale of France during the springtime, suggesting movement during this period (Mignotte et al., 2021). Further illumination of the routes may come from ambitious studies such as MoveInEurope which has a series of radars across the whole of Western Europe, as well as further monitoring of the routes birds take to understand if they are migrating along with the insects to ensure a food source during the journey (Haest, 2024). The routes used by insects and birds in northern Africa may well be linked to those of Western and Eastern Europe.

### Eastern European flyway

The best evidence for the Eastern flyway of Europe is from springtime studies of millions of Diptera (15 families) moving from the Middle East to Cyprus over at least 105 km of ocean (Hawkes et al., 2022). Many bird species have been found to use this route too, a large amount migrating from Eastern Africa before following the Middle Eastern coast (Pedersen et al., 2019). It is expected that at least some of the insects are doing the same thing. The linking of the fertile regions of the Middle East by migratory Diptera in the springtime likely has major importance to eastern European countries in terms of nutrient and pollen transfer (Doyle et al., 2020; Hawkes et al., 2022; Satterfield et al., 2020). During the Autumn season many migratory birds are known to utilise the Georgian corridor to migrate southwards (Verhelst et al., 2011). The Georgian corridor area is difficult to study in terms of Dipteran movements as there is little channelling to ensure the flies move low enough to be counted from ground level, but often insect flyways mirror those of the birds suggesting it is a location worthy of further study.

### Himalayan flyway

The areas north of India and the Himalayas such as Siberia, Mongolia, western China, and Kazakhstan are extensively fertile, but only seasonally during the summer months (Shpedt et al., 2019). Therefore, these are locations migratory Diptera can use to exploit seasonal resources before returning to the fertile lands of the Indian subcontinent during the winter months. Isotopic studies from dragonflies captured in the Maldives suggest that their origins were from southern Siberia, suggesting huge distances are covered by migratory insects using this flyway (Hobson et al., 2012). The great geographic barrier of the Himalayas creates migratory hotspots as the Diptera are directed through mountain passes because of the winds and topography. Therefore, identification of these mountain pass hotspots will allow for easier monitoring of migratory behaviour. A few have been identified but not systematically sampled, providing only tempting morsels of evidence of a long-distance movement of migratory Diptera. *Episyrphus balteatus* Syrphidae were recorded flying through a Nepalese pass at 3700 metres altitude, while various Syrphidae have been seen migrating through the Thorong La pass at 5416 metres altitude (Gatter, 1980; Westmacott & Williams, 1954). However, while only a handful of studies on migratory Diptera exist in this area, it is expected to be a highly fertile area for future study.

### African movements

Due to the size and considerable variety of habitats within the African continent, there are thought to be a great deal of Dipteran migration routes however little is known about them. The great discovery waiting to be made lies within the Northern half of the continent. A recent study based on the normalised difference vegetation index (NDVI) has shown that most suitable habitat for European-summering painted lady butterflies (*Vanessa cardui*) to overwinter is within the sub-Saharan Sahel region (Hu et al., 2021) while field data and ecological niche modelling indicates the Afrotropical region (Talavera et al., 2023). Migratory Syrphidae have been found crossing the Straits of Gibraltar during the springtime suggesting that insects from Africa do recolonise the Europe on their return migration (Ebejer & Bensusan, 2010). NDVI analysis in the Middle East suggested that the numbers of migratory Diptera, like the painted lady butterflies, are also correlated with increased vegetation growth (Hawkes et al., 2022). Therefore, if the Diptera are indeed like the butterflies, they too may be crossing the Sahara to the more favourable Sahel regions. While, to the best of our knowledge, no direct evidence is available for migratory Diptera moving this far, the Bedouin people living at the Bawiti oasis area of Egypt see large numbers of migratory flies moving south in the autumn and north in the spring each year (Mohammed Khozam, *Pers. Comm.*). South of this area, the Saharan desert continues until the Sahel region of Sudan (the next suitable overwintering habitat for these insects). We suggest that European Dipteran migrants may indeed continue across the Sahara on their spring and autumn migrations, making their journeys even more remarkable, but this requires confirmation.

In West Africa in Mali, a total of 28 families of Diptera including Anthomyiidae and Calliphoridae have been recorded making seasonal back and forth movements at altitudes from 40-290m (Florio et al., 2020). Some of these species are likely long-distance migrants that crossed the Sahara, but as the study was primarily nocturnal (aerial traps were opened from 1700-0730) it is possible that many diurnal Dipteran migrants were missed. Other migration routes in Africa include the annual arrival of Simuliidae flies to the Volta River basin in West Africa from distant source areas with the onset of the migration season (Garms et al., 1979). Wind patterns also move large quantities of mosquitoes around West Africa, with the West African monsoon winds enabling large numbers of Dipteran migrants to exploit the seasonal resources created by the monsoon rains (Dao et al., 2014; Huestis et al., 2019; Parker et al., 2005).

Eastern and southern Africa have even fewer studies than west Africa. However, there is some evidence of Dipteran migrants (Glossinidae) arriving with the rains from long distances in Kenya (Brightwell et al., 1997). This suggests that flies here too are utilising the regular seasonal patterns of monsoon winds to migrate, it is likely that far more yet-to-be-discovered taxa are also using these meteorological conditions to exploit seasonal resources in the region (Funk et al., 2016). Africa is an understudied region in terms of Dipteran migration, but there is little doubt there are many migration routes to be discovered.

### East Asia to SE Asia

Long term studies on Beihuang, a small, isolated island in the Bohai Strait, NE China that included trapping, trajectory analysis, and intrinsic markers, revealed that *Episyrphus balteatus* (Syrphidae) exhibit seasonal back and forth latitudinal movement, passing the island each year on long-distance migration (Jia et al., 2022). Population genetic studies have also revealed that *Eupeodes corollae* (Syrphidae) has little differentiation in its population across the whole of China, suggesting regular long-distance movement to maintain geneflow across the whole geographic area (Liu et al., 2019). Migration to the Japanese islands from the Asian mainland may also be regularly occurring. Reports have been made of groups of *Calliphora nigribarbis* (Calliphoridae) flies arriving to southern Japan from the Korean peninsula, some 300 km to the NW during the Autumn migration season (Kurahashi, 1997). Based on phylogenetic analysis of Japanese Encephalitis Virus (JEV) strains found in Japan, it has been determined that at least some of the strains originate in Vietnam and China’s inland region, while others originated in Shanghai, China (Nabeshima et al., 2009). It has been suggested that the mosquito vectors of the disease migrate to the area regularly, brought from SE Asia by a seasonal low level jet stream during the rainy season (which also brings the brown leafhopper (*Laodelphax striatellus* - Hemiptera) to Japan) and on westerly winds from mainland China (Nabeshima et al., 2009).

### Oceania

Like many areas, Australian migratory Diptera are poorly studied. A flyway of various species seems to exist between SE Asia and Northern Australia, especially between Papua New Guinea and Queensland across the Torres Strait. Mosquitoes are thought to enable the regular occurrence of Japanese Encephalitis Virus into Australia from Papua New Guinea, utilising favourable winds (Ritchie & Rochester, 2001). Similar movements are known by the *Culicoides* sp. (Ceratopogonidae) as vectors of diseases including Blue Tongue between Indonesia, Papua New Guinea and Queensland (Eagles et al., 2014). Additionally, movements of *Melangyna* sp. (Syrphidae) have been recorded across the Bass Strait between Tasmania and mainland Australia during the springtime (Hill, 2013), although given the size and climatic variability of the Australian continent, these SE Asia-Australian and Australian-Tasmanian flyways are unlikely to be linked. A citizen science study based on Syrphidae in Australia showed that there were major latitudinal movements throughout the year in four species (*Melangyna viridiceps, Simosyrphus grandicornis, Eristalinus punctulatus*, and *Eristalis tenax*), a behaviour suggestive of migration (Finch & Cook, 2020) however, further work is needed in Australia to reveal the true geographical range of movements of migratory Diptera. *Eristalis tenax* is a cosmopolitan species and appears in migration studies from Europe and North America and is found in Australia and New Zealand, this is also the case for *Episyrphus balteatus* in Europe and East Asia (Finch & Cook, 2020; Hawkes et al., 2022; Jia et al., 2022; Shannon, 1926). The cosmopolitan distribution of these migrants could allow for fascinating studies into the behaviour and genomics of the same species across multiple continents.

### Potential flyways

Vast swathes of the globe are understudied in terms of migratory Diptera and there are undoubtedly more species and flyways to be discovered (as evident from the map in Figure 3). No records of Dipteran migration have been found from sub-equatorial South America. Given that the vast latitudinal difference covered by the landmass will give rise to many seasonal resources to exploit, conditions seem perfect for the presence of migratory Diptera. Similarly, southern Africa is understudied yet has great potential for discovery. One method for discovering new migratory flyways of Diptera is monitoring the routes of migratory birds or the systematic monitoring of insects at likely visible migration points in the landscape. For example, migratory globe-skimmer dragonflies (*Pantala flavescens*) migrate between India and Africa on monsoon winds (R. C. Anderson, 2009) and so smaller Dipteran species may also be traversing the same immense distance to exploit the seasonally available conditions created by the monsoons. Genetic studies have revealed that species of Drosophilidae and Tephritidae found in East Africa have their origins in India, likely having been blown across on the seasonal winds (Jacquard et al., 2013; Tsacas, 1984). Additionally, large numbers of *Chrysomya megacephala* (Calliphoridae) were recorded arriving to a Maldivian island suggesting a similar journey to the *Pantala flavescens* dragonflies (WLH pers. obs.). Thrillingly, also on the Maldives, parasitic *Forcipomyia* midges were recorded clinging to the wings of migratory *Pantala flavescens* dragonflies which had presumably just arrived from India (WLH pers. obs), an example of phoretic migration by these dragon-riding flies.

These tidbits of information on migratory Diptera in this area suggest an important research field for future studies.

## Ecological Roles

We identified a diverse range of Diptera migrants, and they play an equally diverse range of ecological roles. Analysis of the 622 identified species (which often had multiple ecological roles) revealed that 62% were pollinators, 35% were decomposers, 18% were pests, 16% were disease vectors, 10% controlled the pests, and all played a role in the transfer of nutrients (Figure 4). Understanding the roles these Diptera play is imperative when considering how the planet may be impacted by these movements of flies globally.

**Figure 4.**
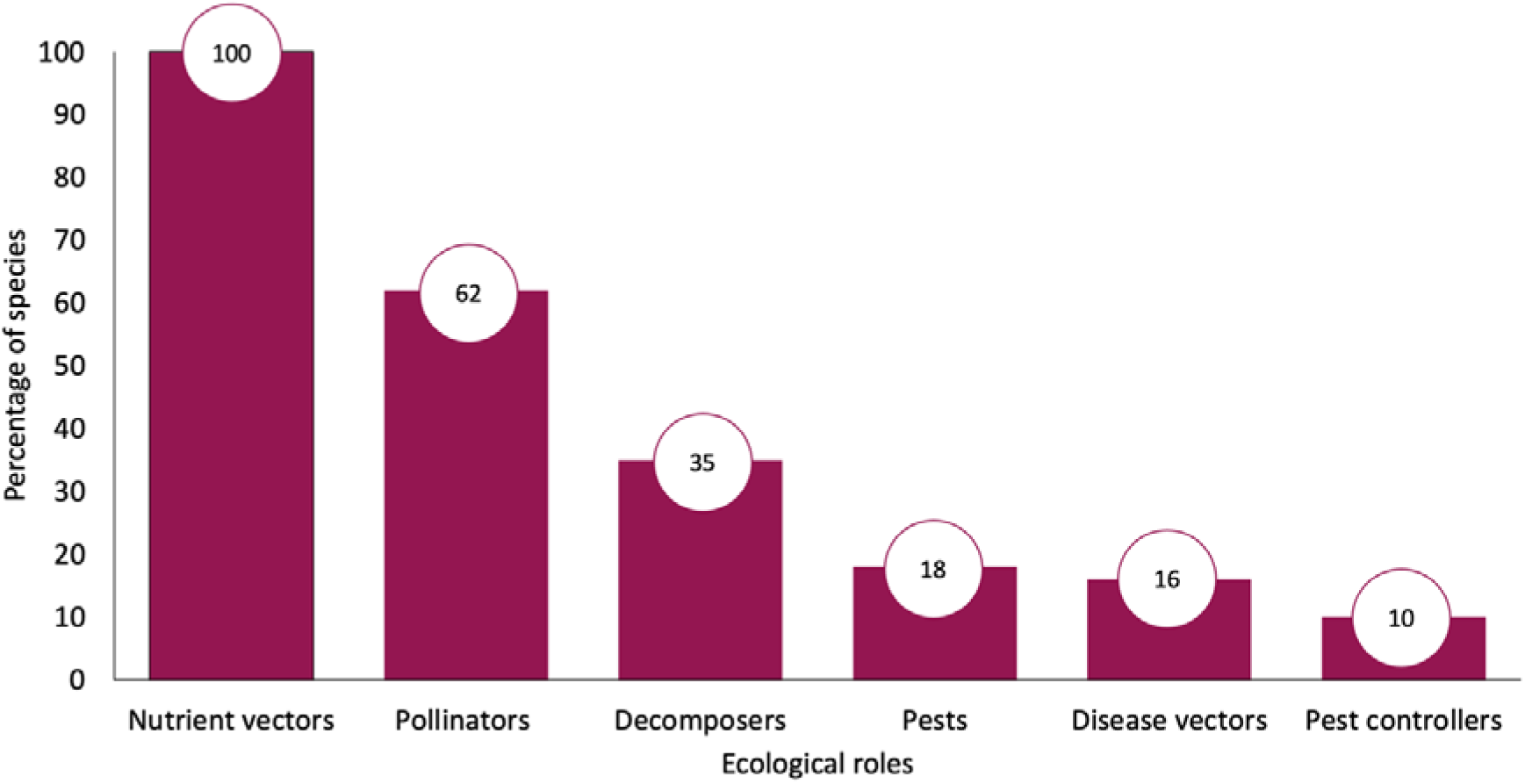
The ecological roles played by the 622 known Dipteran migrants (a single species often played multiple ecological roles).

### Pollinators

An estimated 62% of identified Dipteran migrants have been identified as pollinators. Rader (2020) reviewed non-bee insects as pollinators of crops and found that Diptera visited 72% of major food crops. Six families of flies visited more than 12 major food crops, Syrphidae, Calliphoridae, Muscidae, Sarcophagidae, Tachinidae, and Bombyliidae, and all include species that are known migrants. It was found that amongst these families, the Syrphidae and the Calliphoridae were the most common visitors (Rader et al., 2020). The Syrphidae alone have been found to pollinate 52% of major food crop plants globally with an estimate worth of around US$300 billion per year (Doyle et al., 2020; Rader et al., 2020).

Migratory pollinators may be exceptionally important to global ecosystems because, unlike more sedentary pollinator species, they transport pollen great distances and can link geographically isolated plant populations (Doyle et al., 2020; Lysenkov, 2009; B. Meyer et al., 2009; Rader et al., 2011). Evidence for long-distance transfer of pollen was found for individual *E. tenax* (Syrphidae) and *C. vicina* (Calliphoridae), caught after flying at least 105 km across the eastern Mediterranean from the Middle East to Cyprus with Bug Orchid (*Anacamptis coriophora*) pollen attached to their faces (Hawkes et al., 2022). Further pollen analysis by DNA barcoding revealed that these same *E. tenax* flies were carrying at least seven other species of pollen upon their bodies (Hawkes and T. Doyle *unpublished data*) while data from migratory *E. balteatus* and *E. corollae* caught in the Alps revealed average pollen loads of 10.5 grains per fly (range: 0–107) from up to 3 plant species (Wotton et al. 2019). Pollen can remain viable for up to 2 days (Gibernau et al., 2003) and these insects are capable of moving 100s of km in a matter of hours with wind assistance (Hawkes et al., 2022) suggesting viable pollen can be transferred great distances.

In addition, migratory pollinators may be very numerous; just two species of Syrphidae can transport an estimated 3–8 billion pollen grains into southern Britain from the near continent each year, and 3–19 billion pollen grains out to the continent in the Autumn (Wotton et al. 2019). Such movements are likely to have highly significant consequences for long-range gene flow mediated by insect migration. For example, the movement of pollen may allow for increased gene flow between populations which in turn may increase the resistance of the plants to inbreeding depression, increase the likelihood of the plants surviving and maintain the health of the isolated populations (Luo et al., 2019; Pérez-Bañón et al., 2003). Migratory pollinators may also allow for adaptions by plant populations to counter a warming climate by spreading alleles favourable for disease resistance or drought (Luo et al., 2019; Pérez-Bañón et al., 2003). Small islands without the means to support populations of sedentary pollinators may especially benefit from migratory Dipteran pollinators. For example, in the Columbretes archipelago of Spain the migratory Syrphid *E. tenax* is known to be the major pollinator species, alongside the Calliphorid *Lucillia sericata* (Pérez-Bañón et al., 2007).

### Decomposers

Migratory animals rely on arriving in an area where resources are present upon which their young can develop (Dingle, 2014). Many species of migratory Diptera are decomposers (such as some Calliphoridae and Eristaline Syrphidae), taking the organic matter from a dead organism or an organism’s waste and breaking it down into simple organic substances which can subsequently be taken up by other organisms (Losey & Vaughan, 2006). Studies performed by Hawkes et al (2022) in Cyprus and the Pyrenees (Hawkes et al., 2024) revealed that migratory Diptera, whose life histories play a major role in decomposition, comprise a significant part of the entire migratory assemblage (16% in Cyprus, 33.6% in Pyrenees) (Hawkes et al., 2022; Hawkes et al., 2024) We calculate here that of all known migrant Diptera, those who play a role in decomposition comprise 35%.

Many of the migrant decomposers feed on decaying plant or fungi matter or animal waste. Migratory Calliphoridae such as *L. sericata* (Diakova et al., 2018) and the *C. vicina* group lay their eggs on carrion for their offspring to develop upon (G. S. Anderson, 2011). These carrion feeders are known to fly great distances (Hawkes et al., 2022) and some of their populations are considered panmictic, suggesting high levels of migration (Diakova et al., 2018), therefore the nutrients taken from the larvae feeding on carrion are redistributed across large areas through nutrient transfer. The Syrphidae within the Subfamily Eristalinae are important examples of this. *Eristalis tenax* larvae are coprophagous, saprophagous and aquatic filter feeders which prefer to live in areas with high microbial and organic contamination (Francuski et al., 2014). They therefore aid the biodegradation of organic waste, especially within synanthropic conditions (in association with and benefitting from human activities) (Francuski et al., 2014). In ideal conditions, it has been found that just 8,800 *E. tenax* eggs (0.8ml) (Čičková et al., 2012) can decompose 100 kg of pig slurry, transforming it into organic compost with excellent agronomic potential (Ecodiptera, 2009). This makes this species of Syrphid highly efficient and important decomposers. The impacts migratory Diptera have on decomposition efforts globally are not known, but given their abundance in migratory assemblages, the impacts could be large. Strategies involving the planting of wildflowers and providing other habitats for migratory Diptera near areas where decomposition is needed (livestock slurry pits for example) could allow the maximisation of the decompositional roles of migratory Diptera.

### Pests

The planet has become increasingly agricultural and there are vast swathes of land dedicated to growing the same types of crop or livestock globally (W. B. Meyer & Turner, 1992). Migratory species need to be able to find resources wherever they choose to settle, and the species that have evolved to use these abundant crops or livestock as a food source have been the most successful (Guo et al., 2020). Indeed, monocultures of crops have led to a simplification of the biodiversity of insects, reducing the natural enemies of insects, in turn creating conditions which are suitable for agricultural pests to flourish (Sánchez-Bayo & Wyckhuys, 2019). Many migratory Diptera are classed as agricultural pests, we estimate this number at 18% of all known Dipteran migrants. For example, some migratory species of Chloropidae, such as *Oscinella frit*, are pests of various cereals, grasses, and spring sown maize (El-Wakeil & Volkmar, 2011; Southwood & Jepson, 1962). Over 15 million *Delia platura* (Anthomyiidae) were found migrating long distances (minimum 105 km) from the Middle East to Cyprus in spring 2019, this species is a generalist crop pest of nearly 50 plant species (Guerra et al., 2017; Hawkes et al., 2022). This was the first instance that this species had been recorded migrating in such large numbers, suggesting an increase in either the abundance of this species or the prevalence of its migratory behaviour. Species such as the stable fly *S. calicitrans* (Muscidae) are known costly pests of livestock (particularly cattle) (Campbell et al., 2002; Gerry, 2007), the adult flies feeding on the blood of the mammals to provide a protein source before laying their eggs (Bishopp, 1913). These flies are known to seasonally recolonise dairy farms (Beresford and Sutcliffe, 2009) and can fly at least 225 km based on mark release recapture experiments (Hogsette & Ruff, 1985). *Cochliomyia homininvorax* (Calliphoridae) is a well-known migrant that is a major pest of livestock as its larvae cause myiasis, burrowing into the flesh of the mammal to feed and develop (Costa-Júnior et al., 2019). Methods for controlling these species often include use of pesticides, however rates of pesticide resistance in migratory organisms have been found to be high (Hemingway et al., 1997; M. Raymond & Pasteur, 1996) underlining the need for a greater understanding of the life histories and movement patterns of these species.

### Disease vectors

One of the most important impacts that migratory Diptera have is as vectors of disease, with 16% of identified Dipteran migrants thought to play this role. Of all the migratory Dipteran families, the mosquitoes (Culicidae) which have been the best studied in this regard. Mosquitoes are known vectors of diseases and kill over half a million people globally every year (Bueno-Marí et al., 2022). *Anopheles coluzzii* mosquitoes which are the primary malaria vector have been shown to engage in windborne migration above Africa, travelling up to 300km in 9 hours (Huestis et al., 2019). Of other families, the blackfly *Simulium damosum* (Simuliidae) is capable of moving hundreds of kilometres each year on monsoon winds across west Africa, spreading a nematode (*Onchocerca volvulus*) that causes river blindness (R. H. A. Baker et al., 1990), and *Culicoides* sp. (Ceratopgonidae) are known to aid the seasonal recurrence of blue tongue disease in Israel each year (Braverman & Chechik, 1993). Within the Muscidae, the stable fly *S. calcitrans* is thought to be able to transfer food-associated human pathogens from agricultural to urban areas (Mramba et al., 2007), as well as directly transmitting wildlife diseases (Mihok & Clausen, 1996). Some migratory Diptera are involved in the transmission of plant diseases. For example, the bean seed fly *D. platura* (Anthomyiidae) has been recently discovered as a major vector of the soft rot bacteria (Pasanen, 2020). Another understudied area of research is the role that migratory Syrphidae may play in the transfer of diseases that affect honeybees (*Apis mellifera*) and other bee species, such as deformed wing virus to previously unaffected populations (Fischer et al., 2006). However, although presence of the diseases has been found within the migratory *E. tenax*, there was no evidence of viral replication within the species or data on whether the diseases can be actively passed on to the bees (Fischer et al., 2006).

Mosquito based diseases are generally a major problem within warmer, more tropical, areas of the globe where the Diptera involved in vectoring these diseases (e.g., mosquitoes, Tsetse flies, Simulliidae) can occur in abundance (Huestis et al., 2019). However, with global warming, it is predicted that 4.7 billion more people will be affected by these diseases by 2070 compared to the 1999 numbers (Colón-González et al., 2021). This will be the result of rising temperatures increasing the suitability of locations for the survival of disease vectors. The range expansion of these diseases could cause serious problems in areas where the human population are immunologically naïve, or healthcare systems are unprepared (Colón-González et al., 2021). The migratory behaviour of these Dipteran vectors increases the complexity of combatting the diseases as a new influx of migratory pathogens are introduced each year with the insects’ arrival (e.g., Lebl et al., 2015; Riad et al., 2017). This necessitates the development of management plans which consider the long-distance movement of the vectors. Unfortunately, many insect vectors of disease are understudied in terms of their migratory behaviour, yet targeted research on their movement patterns could have significant impacts for human health.

### Pest controllers

Many arthropods are pests that cause damage to agricultural crops, and many migratory Diptera are predators upon these pests at some stage in their life histories (Courtney et al., 2009). We found that 10% of all migratory Diptera play the role of pest controllers. These pest controllers include many representatives from the Syrphidae (such as the aphidophagous *E. balteatus* and *E. corollae* (Wotton et al., 2019)) as well as from the Calliphoridae (such as *Stomorhina lunata* which feeds upon locust larvae) (Greathead, 1962). Tachinidae are also known to be useful pest controllers as they lay eggs in a variety of insect larvae including those of the Lepidoptera, Coeleoptera, Hemiptera, and Symphyta. For example, the migratory *Tachina fera* (Tachinidae) has been used to control the populations of the Gypsy moth *Lymantria dispar* in forest environments (Davis, 2013). It is presumed that most migrant species are generalists or at least target a highly abundant prey source. As a result of the increased agricultural land coverage, the species that are classed as pests have generally become more dominant in recent times (Guo et al., 2020). Because of this, the migratory Diptera that prey on these pests are increasingly important, especially given the rise of pesticide resistance in populations and the other ecological benefits that migratory Diptera bring to agricultural landscapes (Doyle et al., 2020; Hemingway et al., 1997; M. Raymond & Pasteur, 1996).

Of the migratory Diptera, Syrphidae are best studied regarding pest control (Rojo et al., 2003). Aphidophagous Syrphidae are common migrants across the world’s continents barring Antarctica, meaning the total impact in terms of pest control by the migratory Syrphidae is likely to be huge. Many migratory species such as the abundant *E. balteatus* and *E. corrolae* feed on aphids as larvae and therefore are beneficial to agricultural practices. Indeed, both species are available commercially as biological control agents for use in glasshouses (Moerkens et al., 2021; Pineda & Marcos-García, 2008). The larvae of *E. balteatus* and *E. corrolae* are voracious predators and it has been estimated that the progeny of flies migrating to southern England during the springtime consume up to 10 trillion aphids each year (Wotton et al., 2019). However, the contribution to biological control by migratory Syrphidae is likely to be much greater, as the impact of other immigrations or generations produced by other migratory species is yet to be calculated.

### Nutrient transfer

It is thought that insect migration represents the most important animal movement annually in terrestrial ecosystems, comparable to the most significant of marine migrations (Hu et al., 2016). As migratory Diptera are multi-generational migrants, when they reach a suitable area, they lay their eggs and die (Chapman et al., 2015), hence, 100% of these insects are capable of transporting nutrients between geographically distant ecosystems via carcass deposition (Hu et al., 2016; Satterfield et al., 2020). The dry body weight of a migratory fly is typically comprised of 10% Nitrogen and 1% Phosphorous, elements which are limiting to plant growth (Elser et al., 2000). Therefore, these insects represent a rich source of nutrient influx for ecosystems. Few studies have documented the influence of nutrient transfer by migratory insects, and fewer still have focussed on the Diptera alone. Wotton (Wotton et al., 2019) estimated that the 4 billion *E. balteatus* and *E. corollae* Syrphidae migrating above southern England each year, comprise 80 tons of biomass and will deposit 2500kg of Nitrogen and 250kg of Phosphorous a considerable distance from their source. The entire migratory assemblage moving annually across southern England has been estimated at 3200 tons, 7.7 times the 415 tons of biomass of migrating songbirds, highlighting the huge importance of migratory insects to nutrient transfer (Hu et al., 2016). Migratory Diptera are known to be abundant in migratory assemblages, and by extrapolating the values calculated for the Syrphidae to all other migrant Diptera moving above southern England and the rest of the world, the movement of nutrients each year is likely to be immense.

Far more research is needed into this fascinating field, particularly in high latitude environments, where very few organisms can survive the winter months and where the annual, dependable influx of migratory Diptera into these regions may provide vital nutrients to the continual growth and blooming of vegetation in the area. Animals further up the trophic level which rely on insects as food, such as birds (Tallamy & Shriver, 2021), may rely on the influx of migratory Diptera each year during the springtime to provide the food needed to feed their young. Finally, because migratory Diptera are not aiming for a specific location on their seasonal migrations, it could be that a large percentage of their populations regularly end up drowning in the sea. Migrating Diptera are often trapped on ships far out in the ocean. For example, *Calliphora nigribarbis* and *Aldrichina grahami* (Calliphoridae) were caught 300-450km off Japan in the Pacific Ocean (Kurahashi, 1991). While some flies may eventually reach shore, many more likely drown in the ocean due to exhaustion or inclement weather conditions. In 2022, large numbers of Syrphidae were found stranded on a beach in southwestern France after being caught in a storm and drowning (Fisler & Marcacci, 2022). It could be that these perished flies provide additional nutrients for marine organisms.

## Declines in migratory Diptera

The natural world is under intense pressure from myriad anthropogenically induced impacts, and many insect taxa have undergone precipitous declines: A study monitoring flying insect biomass in Germany revealed a 76% decline over just 27 years (Hallmann et al., 2021). Similarly, in the UK the number of insects found splattered on car numberplates has reduced by 64% in the 18 years since 2004 (Ball et al., 2022). When compared to their sedentary counterparts, however, migratory Diptera that have wide habitat ranges and multiple generations throughout the year, are thought to be more resilient to the effects of climate change (Biesmeijer et al., 2006). Even so, the few studies that do exist on migratory Dipteran declines are still damning. For example, in the last 50 years the number of aphidophagous Syrphidae autumnally migrating through Randecker Maar in the Schwäbische Alb uplands of southwest Germany has declined by 97% (Gatter et al., 2020). These declines may have drastic impacts on the rest of the natural world. For example, North American insectivorous bird numbers have dropped by an estimated 2.9 billion in the last 50 years, compared to non-insectivorous birds whose numbers have increased by 26.2 million individuals (Tallamy & Shriver, 2021). A recent European study on Syrphidae has predicted the loss of some sedentary species from lowland areas and gains in alpine locations (Miličić et al., 2018). The majority of agriculture is found in lowland regions and so the loss of these insect pollinators could negatively impact the crop yield. Migratory species of Syrphidae have high reproductive rates and mobility and, like other insect migrants (M. B. Baker et al., 2015; Bale & Hayward, 2010; Zeng et al., 2020), could be more capable of adapting to climate change, making them particularly important for counteracting damage to the crops caused by poleward shifts in pests such as aphids (Bebber et al., 2013).

These declines are due to a variety of factors, but the main causes are climate change and habitat loss (Goulson, 2019). Insects are thought to be particularly susceptible to extreme weather events such as prolonged droughts or reduced periods of sunshine and increased rainfall (Dennis & Sparks, 2007; Ewald et al., 2015). Agricultural intensification has led to large-scale habitat loss for migratory insects due to monotypic crops and increased pesticide usage preventing the insects from finding a suitable food source while on migration (Benton et al., 2002; Ewald et al., 2015; Grüebler et al., 2008).

Shifts in migratory insect assemblages are expected to have occurred over the last century, with a favouring towards the pest species which rely on human crops. This has meant that the overall insect biomass in many locations has not changed, yet the types of insects of which the biomass is comprised has shifted (Guo et al., 2020). A 15 year systematic monitoring of migratory insects in northeastern Asia showed that while 79% of insect population sizes remained stable over the time period, beneficial insects to humans such as the pest controlling Odonata declined by 90%, and population levels of certain crop pests exhibited an upwards trend (Guo et al., 2015). Further long-term monitoring studies have also found little change in overall biomass over their study’s duration (Hu et al., 2016). However, given the documented declines of many beneficial insect species (Gatter et al., 2020; Hallmann et al., 2021), the numbers of these declining insects are likely being replaced by less beneficial taxa. However, some pest species such as the migratory moths (Silver Y *Autographa gamma,* Black Cutworm *Agrotis ipsilon,* and the turnip moth *Agrotis segetum*) have also shown declines, highlighting the need for further research.

## Climate change and Migratory Diptera

The Earth is currently subject to mass climate change because of anthropogenic actions (Ceballos et al., 2015). Migratory Diptera will not be exempt from the effects of climate change and will be affected in myriad ways. Studies specifically focussing on climate change and migratory Diptera are sparse, yet the main trends are likely reflected in other, better studied, migratory insect taxa and so parallels will be drawn throughout this section.

Global temperatures are likely due to rise between 2-4.9°C above pre-industrial levels by 2100 (Raftery et al., 2017). Increasing temperatures could see higher latitude countries receiving more Dipteran migrants. Correlations between 113 years of migratory Lepidoptera abundance and temperature have shown that these migrants have become more abundant in the UK with increasing temperatures (T. H. Sparks et al., 2007). This is thought to be in part because of increased desiccation in southern Europe encouraging northward migration (T. H. Sparks et al., 2007), something that is likely mirrored by Dipteran migrants. Increasing temperatures in higher latitudes is also increasing the suitability for migrants to persist overwinter. This could lead to the loss or rebalancing of migratory behaviour in many Dipteran species which tend to be partial migrants (Menz, Reynolds, et al., 2019). As a result, the ecological benefits of the Diptera due to their migratory behaviour (as detailed above) will also be lost. Interestingly, the presence of partial migration, where part of the population remains in the breeding area instead of migrating, in many species of migratory insect may lead to a level of resilience to climate shifts (Menz, Reynolds, et al., 2019). For example, some individuals of the migratory Syrphid *E. balteatus* overwinter in parts of central Europe (Luder et al., 2018; Odermatt et al., 2017; L. Raymond et al., 2014), and can do so in all life stages, from eggs, larvae, pupae and adults (L. Raymond et al., 2014). These overwintering animals provide critical early-season control of aphids colonising crops early in the growing season before the migratory individuals have arrived (Raymond et al. 2014). With warming climates, we may see an increase in the proportion of individuals and species overwintering and forgoing migration.

Increasing temperatures due to climate breakdown could also lead to phenological asynchronies between taxa. The timing of Dipteran migration may be linked to temperature as seen in some migratory butterflies such as the red admiral (*Vanessa atalanta*) (T. H. Sparks et al., 2005), indeed the phenology of first sighting of Syrphidae in the UK has advanced earlier in the year as the planet warms (Hassall, et al., 2017). Myriad other organisms may rely upon (or are relied upon by) the arrival of Dipteran migrants such as Passerine birds, which may need the influx of migratory insects to help feed their young, or wildflowers who provide a vital food source for the migrating Diptera (and who may rely on the Diptera for pollination services) (Hawkes et al., 2022; Losey & Vaughan, 2006). If these organisms rely upon day-length and not temperature to dictate their activities, then asynchrony could have disastrous impacts (Mayor et al., 2017). A literature review on the ecological impacts of temperature-mediated trophic asynchrony revealed that there is a dearth in studies on the subject (Samplonius et al., 2021). The studies that do exist are biased towards terrestrial higher trophic secondary consumer taxa such as the birds, and the southern hemisphere is largely understudied (Samplonius et al., 2021). Far more research is needed in this field to inform conservation efforts and to understand the possible consequences of phenological change.

The range of many wind-borne Dipteran migrants are expanding in response to increases in temperature, as seen in some *Aedes* spp. Mosquitoes, which are important vectors of diseases such as malaria. Wind patterns that bore mosquitoes to high altitude settlements in the Himalayan region used to pose no threat to the humans as the cold temperatures would kill the mosquitoes (Dhimal et al., 2021). However, with global warming the mosquitoes can now survive in these regions and transmit fatal diseases such as malaria, thus posing a serious threat to these unprepared communities (Dhimal et al., 2021). This problem is not limited to this one example, as increasing temperatures globally mean that higher latitude countries are now at threat from these mosquito-vectored diseases due to the increased favourability in conditions for mosquito survival (Agyekum et al., 2021). Furthermore, the response of disease vectors to changes in climatic conditions within their distribution has significant consequences for predicting and managing outbreaks of disease.

Increased extreme weather events such as long droughts or extended periods of rainfall due to climate breakdown are thought to have negative impacts on migratory Diptera populations due to changes in habitat suitability. Increased drought may cause vegetation to wither prematurely and the eggs of Diptera to dry out and become unviable. Similarly, increased rainfall may be detrimental to Diptera larvae that develop underground, as they can drown in the waterlogged soil. However, droughts and the loss of moist habitats can lead to a reduction in the availability of suitable breeding sites for many saprophagous species that have semi-aquatic larvae, such as the Syrphid *E. tenax*.

Finally, increased CO_2_ levels have been shown to reduce the amount of nitrogen in plant leaves by 10-30%. As a result, herbivory levels by crop pests (including many migratory Diptera) are thought to increase 20-90% to compensate for this reduced nitrogen availability (Kinney et al., 1997; Roth & Lindroth, 1994, 1995), potentially leading to increased crop damage and resultant costs to growers. Very little is known about the response of migratory Diptera to climate change. Research into the ecological roles, range-shifts, and declines of these hugely important species is desperately needed so that their impacts can be understood and either encouraged or mitigated, particularly in the context of ecosystem and human health.

## Conclusion and future research

Our analyses of the literature on migrant Diptera have revealed a highly diverse set of species, many of which appears to be highly abundant, and to migrate in huge numbers. They carry out a wide range of ecological roles that impact a large swathe of the globe and because of this, they should be considered an important and remarkable group of migrants globally. However, compared to other groups, very little is known and for many of the migratory families only a single study related to migration was uncovered. We recommend a greater focus on the diversity of Dipteran migrants, as many new discoveries await as we see in the new migratory behaviour uncovered in recent studies (Hawkes et al., 2022, 2024 under review). In addition, this will help to understand the ecological roles these insects are performing as they connect distant landscapes. Techniques such as monitoring and trapping in migration hotspots, stable isotope and pollen analysis, trajectory analysis, flight simulators and flight mills, and NDVI measurements, along with emerging approaches such as networks of radar, can be used to infer behaviour, assemblages, origins, destinations and the headings and numbers of mass movements of Dipteran migrants.

Many anthropogenically beneficial Dipteran migrants are under threat from climate change and other anthropogenic impacts. It is possible that many migratory flies and their behaviour could disappear without even being documented. To conserve these vitally important taxa, it is not enough to simply protect or restore habitat at one location: the entire migratory route must be capable of sustaining these insects, as for other migratory species (Runge et al., 2014). Slightly altering agricultural, rewilding and conservation practices to ensure landscape connectivity could have the greatest impact in this regard. Migratory pollinators from other taxa are known to use corridors of sequentially blooming flowers and any loss of flowering plant populations along these corridors could have severe negative impacts on the survival of migratory Diptera (Nabhan, 2004). The maintenance of hedgerows and other woody structures in otherwise barren agricultural landscapes can also provide key microclimate refugia for overwintering or oversummering individuals (Raymond et al. 2014). Reducing pesticide usage and providing wildflower strips alongside (or within) fields for these migratory Diptera to feed on during migration would be of major help (Haaland et al., 2011). In setting future conservation measures, it is key to understand the migratory cycles and pathways of these ecologically important species. It is hoped that this review inspires many further studies into these remarkable Dipteran migrants.

## References

Agyekum, T. P., Botwe, P. K., Arko-Mensah, J., Issah, I., Acquah, A. A., Hogarh, J. N., Dwomoh, D., Robins, T. G., & Fobil, J. N. (2021). A systematic review of the effects of temperature on Anopheles mosquito development and survival: implications for malaria control in a future warmer climate. International Journal of Environmental Research and Public Health, 18(14), 7255.

Anderson, G. S. (2011). Comparison of decomposition rates and faunal colonization of carrion in indoor and outdoor environments. Journal of Forensic Sciences, 56(1), 136–142.

Anderson, R. C. (2009). Do dragonflies migrate across the western Indian Ocean? Journal of Tropical Ecology, 25(4), 347–358.

Ashmole, N.P. and Ashmole, M.J., 1988. Insect dispersal on Tenerife, Canary Islands: high altitude fallout and seaward drift. Arctic and Alpine Research, 20(1), pp.1–12.

Aubert, J., Aubert, J.-J., & Goeldlin, P. (1976). Twelve years of systematic collecting of syrphids (Diptera) at the Bretolet pass (Alps of Valais). Mitteilungen Der Schweizerischen Entomologischen Gesellschaft, 49(1/2), 115–142.

Babic, I., Baranov, N., & Ganslmayer, R. (1935). The Golubatz Fly in 1934. Arch. Tierheilk., 69(3).

Baker, M. B., Venugopal, P. D., & Lamp, W. O. (2015). Climate change and phenology: Empoasca fabae (Hemiptera: Cicadellidae) migration and severity of impact. PLoS One, 10(5), e0124915.

Baker, R. H. A., Guillet, P., Seketeli, A., Poudiougo, P., Boakye, D., Wilson, M. D., & Bissan, Y. (1990). Progress in controlling the reinvasion of windborne vectors into the western area of the Onchocerciasis Control Programme in West Africa. Philosophical Transactions of the Royal Society of London. B, Biological Sciences, 328(1251), 731–750.

Bale, J. S., & Hayward, S. A. L. (2010). Insect overwintering in a changing climate. Journal of Experimental Biology, 213(6), 980–994.

Ball, L., Still, R., Riggs, A., Skilbeck, A., Shardlow, M., Whitehouse, A., & Tinsley-Marshall, P. (2022). The Bugs Matter Citizen Science Survey: Counting insect’splats’ on vehicle number plates. policycommons.net

Bebber, D. P., Ramotowski, M. A. T., & Gurr, S. J. (2013). Crop pests and pathogens move polewards in a warming world. Nature Climate Change, 3(11), 985–988.

Beebe, W. (1951). Migration of insects (other than Lepidoptera) through Portachuelo Pass, Rancho Grande, north-central Venezuela. Zoologica: Scientific Contributions of the New York Zoological Society, 36(20), 255–266.

Benton, T. G., Bryant, D. M., Cole, L., & Crick, H. Q. P. (2002). Linking agricultural practice to insect and bird populations: a historical study over three decades. Journal of Applied Ecology, 39(4), 673–687.

Biesmeijer, J. C., Roberts, S. P. M., Reemer, M., Ohlemuller, R., Edwards, M., Peeters, T., Schaffers, A. P., Potts, S. G., Kleukers, R., & Thomas, C. D. (2006). Parallel declines in pollinators and insect-pollinated plants in Britain and the Netherlands. Science, 313(5785), 351–354.

Bishopp, F. C. (1913). The stable fly (Stomoxys calcitrans L.), an important live stock pest. Journal of Economic Entomology, 6(1), 112–126.

Braverman, I. v, & Chechik, P. (1993). Introduction of Culicoides (Diptera, Ceratopogonidae). Israel Journal of Veterinary Medicine, 48.

Brenton, L. C. L. (1844). The Septuagint Version of the Old Testament, According to the Vatican Text, Tr. Into English: with the Principal Various Readings of the Alexandrine Copy, and a Table of Comparative Chronology (Vol. 1). S. Bagster.

Brightwell, R., Dransfield, R. D., Stevenson, P., & Williams, B. (1997). Changes over twelve years in populations of Glossina pallidipes and Glossina longipennis (Diptera: Glossinidae) subject to varying trapping pressure at Nguruman, south-west Kenya. Bulletin of Entomological Research, 87(4), 349–370.

Bueno-Marí, R., Drago, A., Montalvo, T., Dutto, M., & Becker, N. (2022). Classic and novel tools for mosquito control worldwide. Ecology and Control of Vector-borne Diseases (pp. 234–238). Wageningen Academic Publishers.

Campbell, J. B., Boxler, D. J., & Adams, D. C. (2002). Stable fly, Stomoxys calcitrans, (Diptera: Muscidae) numbers trapped at Nebraska sandhill pasture sites from 1998-2002. The 2002 ESA Annual Meeting and Exhibition.

Ceballos, G., Ehrlich, P. R., Barnosky, A. D., García, A., Pringle, R. M., & Palmer, T. M. (2015). Accelerated modern human–induced species losses: Entering the sixth mass extinction. Science Advances, 1(5), e1400253.

Chapman, J. W., Nesbit, R. L., Burgin, L. E., Reynolds, D. R., Smith, A. D., Middleton, D. R., & Hill, J. K. (2010). Flight orientation behaviors promote optimal migration trajectories in high-flying insects. Science, 327(5966), 682–685.

Chapman, J. W., Nilsson, C., Lim, K. S., Bäckman, J., Reynolds, D. R., & Alerstam, T. (2016). Adaptive strategies in nocturnally migrating insects and songbirds: contrasting responses to wind. Journal of Animal Ecology, 85(1), 115–124.

Chapman, J. W., Reynolds, D. R., Smith, A. D., Smith, E. T., & Woiwod, I. P. (2004). An aerial netting study of insects migrating at high altitude over England. Bulletin of Entomological Research, 94(2). 10.1079/ber2004287

Chapman, J. W., Reynolds, D. R., & Wilson, K. (2015). Long-range seasonal migration in insects: mechanisms, evolutionary drivers and ecological consequences. Ecology Letters, 18(3), 287–302.

Chowdhury, S., Fuller, R.A., Dingle, H., Chapman, J.W. and Zalucki, M.P., 2021. Migration in butterflies: a global overview. Biological Reviews, 96(4), pp.1462–1483.

Chown K, S. L. & L. (1994). Recently established Diptera and Lepidoptera on sub-antarctic Marion Island. African Entomology, 2(1), 57–60.

Čičková, H., Pastor, B., Kozánek, M., Martínez-Sánchez, A., Rojo, S., & Takáč, P. (2012). Biodegradation of pig manure by the housefly, Musca domestica: a viable ecological strategy for pig manure management. Plos One, 7(3), e32798.

Clem, C.S., Hobson, K.A. and Harmon-Threatt, A.N., (2023). Insights into natal origins of migratory Nearctic hover flies (Diptera: Syrphidae): new evidence from stable isotope (δ2H) assignment analyses. Ecography, 2023(2), p.e06465.

Colón-González, F. J., Sewe, M. O., Tompkins, A. M., Sjödin, H., Casallas, A., Rocklöv, J., Caminade, C., & Lowe, R. (2021). Projecting the risk of mosquito-borne diseases in a warmer and more populated world: a multi-model, multi-scenario intercomparison modelling study. The Lancet Planetary Health, 5(7), e404–e414.

Costa-Júnior, L. M., Chaves, D. P., Brito, D. R. B., Santos, V. A. F. dos, Costa-Júnior, H. N., & Barros, A. T. M. (2019). A review on the occurrence of Cochliomyia hominivorax (Diptera: Calliphoridae) in Brazil. Revista Brasileira de Parasitologia Veterinária, 28, 548–562.

Courtney, G. W., Pape, T., Skevington, J. H., & Sinclair, B. J. (2009). Biodiversity of diptera. Insect Biodiversity.

Dao, A., Yaro, A. S., Diallo, M., Timbiné, S., Huestis, D. L., Kassogué, Y., Traoré, A. I., Sanogo, Z. L., Samaké, D., & Lehmann, T. (2014). Signatures of aestivation and migration in Sahelian malaria mosquito populations. Nature, 516(7531), 387–390.

Davis, D. J. (2013). The phylogenetics of Tachinidae (Insecta: Diptera) with an emphasis on subfamily structure. (Doctoral dissertation, Wright State University)

Dennis, R. L. H., & Sparks, T. H. (2007). Climate signals are reflected in an 89 year series of British Lepidoptera records. European Journal of Entomology, 104(4), 763.

Dhimal, M., Kramer, I. M., Phuyal, P., Budhathoki, S. S., Hartke, J., Ahrens, B., Kuch, U., Groneberg, D. A., Nepal, S., & Liu, Q.-Y. (2021). Climate change and its association with the expansion of vectors and vector-borne diseases in the Hindu Kush Himalayan region: a systematic synthesis of the literature. Advances in Climate Change Research, 12(3), 421–429.

Diakova, A. v, Schepetov, D. M., Oyun, N. Y., Shatalkin, A. I., & Galinskaya, T. v. (2018). Assessing genetic and morphological variation in populations of Eastern European Lucilia sericata (Diptera: Calliphoridae). European Journal of Entomology, 115, 192–197.

Dingle, H. (2014). Migration: the biology of life on the move. Oxford University Press, USA.

Doyle, T., Hawkes, W. L. S., Massy, R., Powney, G. D., Menz, M. H. M., & Wotton, K. R. (2020). Pollination by hoverflies in the Anthropocene: Pollination by Hoverflies. In Proceedings of the Royal Society B: Biological Sciences, 287, 1927. 10.1098/rspb.2020.0508

Doyle, T., Jimenez-Guri, E., Hawkes, W. L. S., Massy, R., Mantica, F., Permanyer, J., Cozzuto, L., Hermoso Pulido, T., Baril, T., & Hayward, A. (2022). Genome-wide transcriptomic changes reveal the genetic pathways involved in insect migration. Molecular Ecology, 31(16), 4332–4350.

Eagles, D., Melville, L., Weir, R., Davis, S., Bellis, G., Zalucki, M. P., Walker, P. J., & Durr, P. A. (2014). Long-distance aerial dispersal modelling of Culicoides biting midges: case studies of incursions into Australia. BMC Veterinary Research, 10(1), 1–10.

Ebejer, M. J., & Bensusan, K. (2010). Hoverflies (Diptera, Syrphidae) recently encountered on Gibraltar, with two species new for Iberia. Dipterists Digest, 17, 123–139.

Ecodiptera. (2009). Ecodiptera – implementation of a management model for the ecologically sustainable treatment of pig manure in the Region of Los Serranos, Valencia-Spain. Available at: https://tinyurl.com/y2qg4e3g

Elser, J. J., Fagan, W. F., Denno, R. F., Dobberfuhl, D. R., Folarin, A., Huberty, A., Interlandi, S., Kilham, S. S., McCauley, E., & Schulz, K. L. (2000). Nutritional constraints in terrestrial and freshwater food webs. Nature, 408(6812), 578–580.

El-Wakeil, N., & Volkmar, C. (2011). Effect of weather conditions on frit fly (Oscinella frit, Diptera: Chloropidae) activity and infestation levels in spring wheat in central Germany. Gesunde Pflanzen, 63(4), 159–165.

Ewald, J. A., Wheatley, C. J., Aebischer, N. J., Moreby, S. J., Duffield, S. J., Crick, H. Q. P., & Morecroft, M. B. (2015). Influences of extreme weather, climate and pesticide use on invertebrates in cereal fields over 42 years. Global Change Biology, 21(11), 3931–3950.

Finch, J. T. D., & Cook, J. M. (2020). Flies on vacation: evidence for the migration of Australian Syrphidae (Diptera). Ecological Entomology, 45(4), 896–900.

Fischer, O. A., Matlova, L., Dvorska, L., Švástová, P., Bartoš, M., Weston, R. T., & Pavlik, I. (2006). Various stages in the life cycle of syrphid flies (Eristalis tenax; Diptera: Syrphidae) as potential mechanical vectors of pathogens causing mycobacterial infections in pig herds. Folia Microbiologica, 51(2), 147–153.

Fisler, L., & Marcacci, G. (2022). Tens of thousands of migrating hoverflies found dead on a strandline in the South of France. Insect Conservation and Diversity. DOI: 10.1111/icad.12616

Fleming, T. H., Eby, P., Kunz, T. H., & Fenton, M. B. (2003). Ecology of bat migration. Bat Ecology, 156, 164–165.

Florio, J., Verú, L. M., Dao, A., Yaro, A. S., Diallo, M., Sanogo, Z. L., Samaké, D., Huestis, D. L., Yossi, O., & Talamas, E. (2020). Diversity, dynamics, direction, and magnitude of high-altitude migrating insects in the Sahel. Scientific Reports, 10(1), 1–14.

Francuski, L., Djurakic, M., Ludoški, J., Hurtado, P., Pérez-Bañón, C., Ståhls, G., Rojo, S., & Milankov, V. (2014). Shift in phenotypic variation coupled with rapid loss of genetic diversity in captive populations of Eristalis tenax (Diptera: Syrphidae): consequences for rearing and potential commercial use. Journal of Economic Entomology, 107(2), 821–832.

Funk, C., Hoell, A., Shukla, S., Husak, G., & Michaelsen, J. (2016). The East African monsoon system: seasonal climatologies and recent variations. The Monsoons and Climate Change: Observations and Modeling, 163–185.

Gao, B., Wotton, K. R., Hawkes, W. L. S., Menz, M. H. M., Reynolds, D. R., Zhai, B.-P., Hu, G., & Chapman, J. W. (2020). Adaptive strategies of high-flying migratory hoverflies in response to wind currents. Proceedings of the Royal Society B, 287(1928), 20200406.

Garms, R., Walsh, J. F., & Davies, J. B. (1979). Studies on the reinvasion of the Onchocerciasis Control Programme in the Volta River Basin by Simulium damnosum sI with emphasis on the south-western areas. Tropenmedizin Und Parasitologie, 30(3), 345–362.

Gatter, W., 1977. Eine Wanderung der Erdschnake (Tipula oleracea l.). Passive Verdriftung oder gerichtete Migration. Nachrichtenblatt Bayeri scher Entomologen, 26, pp.141–152.

Gatter, W. (1980). Nordwärts gerichtete Frühjahrswanderungen palaearktischer Schmetterlinge, Fliegen und Hummeln im Himalaya-und Transhimalayagebiet Nepals. Atalanta, 11, 188–196.

Gatter, W., Ebenhöh, H., Kima, R., Gatter, W., & Scherer, F. (2020). 50-jährige Untersuchungen an migrierenden Schwebfliegen, Waffenfliegen und Schlupfwespen belegen extreme Rückgänge (Diptera: Syrphidae, Stratiomyidae; Hymenoptera: Ichneumonidae). Entomologische Zeitschrift Schwanfeld, 130(3), 131–142.

Gerry, A. C. (2007). Predicting and controlling stable flies on California dairies. UCANR Publications.

Gibernau, M., Macquart, D., Diaz, A., House, D., Fern-Barrow, P., & Dorset, B. (2003). Pollen viability and longevity in two species of Arum. Aroideana, 26, 58–62.

Glick, P.A., 1939. The distribution of insects, spiders, and mites in the air. United States Department of Agriculture Technical bulletin 673, 1–151.

Goulson, D. (2019). The insect apocalypse, and why it matters. Current Biology, 29(19), R967– R971.

Greathead, D. J. (1962). The biology of Stomorhina lunata (Fabricius)(Diptera: Calliphoridae), a predator of the eggs of Acrididae. Proceedings of the Zoological Society of London, 139(1), 139–180.

Grüebler, M. U., Morand, M., & Naef-Daenzer, B. (2008). A predictive model of the density of airborne insects in agricultural environments. Agriculture, Ecosystems & Environment, 123(1–3), 75–80.

Guerra, P. C., Keil, C. B., Stevenson, P. C., Mina, D., Samaniego, S., Peralta, E., Mazon, N., & Chancellor, T. C. B. (2017). Larval performance and adult attraction of Delia platura (Diptera: Anthomyiidae) in a native and an introduced crop. Journal of Economic Entomology, 110(1), 186–191.

Guo, J., Fu, X., Wu, X., Zhao, X., & Wu, K. (2015). Annual migration of Agrotis segetum (Lepidoptera: Noctuidae): observed on a small isolated island in northern China. PLoS One, 10(6), e0131639.

Guo, J., Fu, X., Zhao, S., Shen, X., Wyckhuys, K. A. G., & Wu, K. (2020). Long-term shifts in abundance of (migratory) crop-feeding and beneficial insect species in northeastern Asia. Journal of Pest Science, 93(2), 583–594.

Haaland, C., Naisbit, R. E., & Bersier, L. (2011). Sown wildflower strips for insect conservation: a review. Insect Conservation and Diversity, 4(1), 60–80.

Haest B, Liechti F, Hawkes WL, Chapman J, Åkesson S, Shamoun-Baranes J, Nesterova AP, Comor V, Preatoni D, Bauer S. Accepted. Continental-scale patterns in diel flight timing of high-altitude migratory insects. Philosophical Transactions of the Royal Society B

Hallmann, C. A., Ssymank, A., Sorg, M., de Kroon, H., & Jongejans, E. (2021). Insect biomass decline scaled to species diversity: General patterns derived from a hoverfly community. Proceedings of the National Academy of Sciences, 118(2), e2002554117.

Hardy, A. C., & Cheng, L. (1986). Studies in the distribution of insects by aerial currents. III. Insect drift over the sea. Ecological Entomology, 11(3), 283–290.

Hassall, Christopher, Jennifer Owen, and Francis Gilbert. “Phenological shifts in hoverflies (Diptera: Syrphidae): linking measurement and mechanism.” Ecography 40 7 (2017): 853–863.

Hawkes, W. L. S., Walliker, E., Gao, B., Forster, O., Lacey, K., Doyle, T., Massy, R., Roberts, N. W., Reynolds, D. R., & Özden, Ö. (2022). Huge spring migrations of insects from the Middle East to Europe: quantifying the migratory assemblage and ecosystem services. Ecography, e06288.

Hawkes, W. L., Weston, S. T., Cook, H., Doyle, T., Massy, R., Guri, E. J., Wotton Jimenez, R. E., & Wotton, K. R. (2022). Migratory hoverflies orientate north during spring migration. Biology Letters, 18(10), 20220318.

Hemingway, J., Penilla, R. P., Rodriguez, A. D., James, B. M., Edge W and Rogers, H., & Rodriguez, M. H. (1997). Resistance management strategies in malaria vector mosquito control. A large-scale field trial in Southern Mexico. Pesticide Science, 51(3), 375–382.

Hill, L. (2013). Long-term light trap data from Tasmania, Australia. Plant Protection Quarterly, 28(1), 22–27.

Hlaváček, A., Lučan, R. K., & Hadrava, J. (2022). Autumnal migration patterns of hoverflies (Diptera: Syrphidae): interannual variability in timing and sex ratio. PeerJ, 10, e14393.

Hobson, K. A., Anderson, R. C., Soto, D. X., & Wassenaar, L. I. (2012). Isotopic Evidence That Dragonflies (Pantala flavescens) Migrating through the Maldives Come from the Northern Indian Subcontinent. In PLoS ONE 7, 12. 10.1371/journal.pone.0052594

Hogsette, J. A., & Ruff, J. P. (1985). Stable fly (Diptera: Muscidae) migration in northwest Florida. Environmental Entomology, 14(2), 170–175.

Hondelmann, P., & Poehling, H. (2007). Diapause and overwintering of the hoverfly Episyrphus balteatus. Entomologia Experimentalis et Applicata, 124(2), 189–200.

Howard, E., & Davis, A. K. (2009). The fall migration flyways of monarch butterflies in eastern North America revealed by citizen scientists. Journal of Insect Conservation, 13, 279–286.

Hu, G., Lim, K. S., Horvitz, N., Clark, S. J., Reynolds, D. R., Sapir, N., & Chapman, J. W. (2016). Mass seasonal bioflows of high-flying insect migrants. Science 354, 6319, pp. 1584–1587. 10.1126/science.aah4379

Hu, G., Stefanescu, C., Oliver, T. H., Roy, D. B., Brereton, T., van Swaay, C., Reynolds, D. R., & Chapman, J. W. (2021). Environmental drivers of annual population fluctuations in a trans-Saharan insect migrant. Proceedings of the National Academy of Sciences, 118(26).

Huestis, D. L., Dao, A., Diallo, M., Sanogo, Z. L., Samake, D., Yaro, A. S., Ousman, Y., Linton, Y.-M., Krishna, A., & Veru, L. (2019). Windborne long-distance migration of malaria mosquitoes in the Sahel. Nature, 574(7778), 404–408.

Jacquard, C., Virgilio, M., David, P., Quilici, S., de Meyer, M., & Delatte, H. (2013). Population structure of the melon fly, Bactrocera cucurbitae, in Reunion Island. Biological Invasions, 15, 759–773.

Jensen, J.-K. (2001). An invasion of migrating insects (Syrphidae and Lepidoptera) on the Faroe Islands in September 2000. Norwegian Journal of Entomology, 48, 263–268.

Jia, H., Liu, Y., Li, X., Li, H., Pan, Y., Hu, C., Zhou, X., Wyckhuys, K. A. G., & Wu, K. (2022). Windborne migration amplifies insect-mediated pollination services. Elife, 11, e76230.

Jones, C. M., Parry, H., Tay, W. T., Reynolds, D. R., & Chapman, J. W. (2019). Movement ecology of pest Helicoverpa: implications for ongoing spread. Annual Review of Entomology, 64, 277–295.

Keaster, A. J., Grundler, J. A., Craig, W. S., & Jackson, M. A. (1996). Noctuid moths and other insects captured in wing-style traps baited with black cutworm (Lepidoptera: Noctuidae) pheromone on offshore oil platforms in the Gulf of Mexico, 1988-1991. Journal of the Kansas Entomological Society, 17–25.

Kennedy, J. S. (1985). Migration, behavioral and ecological. Migration: Mechanisms and Adaptive Significance.

Kinney, K. K., Lindroth, R. L., Jung, S. M., & Nordheim, E. v. (1997). Effects of CO2 and NO3−availability on deciduous trees: phytochemistry and insect performance. Ecology, 78(1), 215–230.

Knoblauch, A., Thoma, M., & Menz, M. H. M. (2021). Autumn southward migration of dragonflies along the Baltic coast and the influence of weather on flight behaviour. Animal Behaviour, 176, 99–109.

Kurahashi, H. (1991). The calyptrate muscoid flies collected on weather ships located at the ocean weather stations. Japanese Journal of Sanitary Zoology, 42(1), 53–55.

Kurahashi, H. (1997). Witnessing hundreds of Calliphora nigribarbis in migratory flight and landing in Nagasaki, Western Japan. Med. Entomol. Zool., 48, 55–58.

Lack, D., & Lack, E. (1951). Migration of insects and birds through a Pyrenean pass. The Journal of Animal Ecology, 63–67.

Lebl, K., Zittra, C., Silbermayr, K., Obwaller, A., Berer, D., Brugger, K., Walter, M., Pinior, B., Fuehrer, H.-P., & Rubel, F. (2015). Mosquitoes (Diptera: Culicidae) and their relevance as disease vectors in the city of Vienna, Austria. Parasitology Research, 114, 707–713.

Li, X., Wu, M., Ma, J., Gao, B., Wu, Q., Chen, A., Liu, J., Jiang, Y., Zhai, B., & Early, R. (2020). Prediction of migratory routes of the invasive fall armyworm in eastern China using a trajectory analytical approach. Pest Management Science, 76(2), 454–463.

Liu, M., Wang, X., Ma, L., Cao, L., Liu, H., Pu, D., & Wei, S. (2019). Genome-wide developed microsatellites reveal a weak population differentiation in the hoverfly Eupeodes corollae (Diptera: Syrphidae) across China. Plos One, 14(9), e0215888.

Losey, J. E., & Vaughan, M. (2006). The economic value of ecological services provided by insects. Bioscience, 56(4), 311–323.

Luder, K., Knop, E., & Menz, M. H. M. (2018). Contrasting responses in community structure and phenology of migratory and non-migratory pollinators to urbanization. Diversity and Distributions, 24(7), 919–927.

Luo, L., Xia, H., & Lu, B.-R. (2019). Crop breeding for drought resistance. Frontiers in Plant Science, 10, 314.

Lysenkov, S. N. (2009). On the estimation of the influence of the character of insect pollinators movements on the pollen transfer dynamics. Entomological Review, 89(2), 143–149.

Massy, R., Hawkes, W. L. S., Doyle, T., Troscianko, J., Menz, M. H. M., Roberts, N. W., Chapman, J. W., & Wotton, K. R. (2021). Hoverflies use a time-compensated sun compass to orientate during autumn migration. Proceedings of the Royal Society B, 288(1959), 20211805.

Mayor, S. J., Guralnick, R. P., Tingley, M. W., Otegui, J., Withey, J. C., Elmendorf, S. C., Andrew, M. E., Leyk, S., Pearse, I. S., & Schneider, D. C. (2017). Increasing phenological asynchrony between spring green-up and arrival of migratory birds. Scientific Reports, 7(1), 1902.

Menz, M. H. M., Brown, B. v., & Wotton, K. R. (2019). Quantification of migrant hoverfly movements (diptera: syrphidae) on the west coast of North America. In Royal Society Open Science 6, 4. 10.1098/rsos.190153

Menz, M. H. M., Reynolds, D. R., Gao, B., Hu, G., Chapman, J. W., & Wotton, K. R. (2019). Mechanisms and consequences of partial migration in insects. Frontiers in Ecology and Evolution, 7, 403.

Menz, M. H. M., Scacco, M., Bürki-Spycher, H.-M., Williams, H. J., Reynolds, D. R., Chapman, J. W., & Wikelski, M. (2022). Individual tracking reveals long-distance flight-path control in a nocturnally migrating moth. Science, 377(6607), 764–768.

Meyer, B., Jauker, F., & Steffan-Dewenter, I. (2009). Contrasting resource-dependent responses of hoverfly richness and density to landscape structure. Basic and Applied Ecology, 10(2), 178–186.

Meyer, W. B., & Turner, B. L. (1992). Human population growth and global land-use/cover change. Annual Review of Ecology and Systematics, 39–61.

Mignotte, A., Garros, C., Dellicour, S., Jacquot, M., Gilbert, M., Gardès, L., Balenghien, T., Duhayon, M., Rakotoarivony, I., de Wavrechin, M. and Huber, K., 2021. High dispersal capacity of Culicoides obsoletus (Diptera: Ceratopogonidae), vector of bluetongue and Schmallenberg viruses, revealed by landscape genetic analyses. Parasites & Vectors, 14(1), pp.1–14.

Mihok, S., & Clausen, P. H. (1996). Feeding habits of Stomoxys spp. stable flies in a Kenyan forest. Medical and Veterinary Entomology, 10(4), 392–394.

Miličić, M., Vujić, A., & Cardoso, P. (2018). Effects of climate change on the distribution of hoverfly species (Diptera: Syrphidae) in Southeast Europe. Biodiversity and Conservation, 27(5), 1173–1187.

Moerkens, R., Boonen, S., Wäckers, F. L., & Pekas, A. (2021). Aphidophagous hoverflies reduce foxglove aphid infestations and improve seed set and fruit yield in sweet pepper. Pest Management Science, 77(6), 2690–2696.

Mramba, F., Broce, A. B., & Zurek, L. (2007). Vector competence of stable flies, Stomoxys calcitrans L.(Diptera: Muscidae), for Enterobacter sakazakii. Journal of Vector Ecology, 32(1), 134–139.

Nabeshima, T., Loan, H. T. K., Inoue, S., Sumiyoshi, M., Haruta, Y., Nga, P. T., Huoung, V. T. Q., del Carmen Parquet, M., Hasebe, F., & Morita, K. (2009). Evidence of frequent introductions of Japanese encephalitis virus from south-east Asia and continental east Asia to Japan. Journal of General Virology, 90(4), 827–832.

Nabhan, G. P. (2004). Conserving migratory pollinators and nectar corridors in western North America. University of Arizona Press.

Nielsen, T. R., Andreassen, A. T., & Leendertse A, S. S. (2010). A migration of the Hoverfly Helophilus trivittatus (Fabricius, 1805)(Diptera, Syrphidae) to SW Norway in 2010. Norwegian Journal of Entomology, 57, 136–138.

Nilssen, A. C., & Anderson, J. R. (1995). Flight capacity of the reindeer warble fly, Hypoderma tarandi (L.), and the reindeer nose bot fly, Cephenemyia trompe (Modeer)(Diptera: Oestridae). Canadian Journal of Zoology, 73(7), 1228–1238.

Odermatt, J., Frommen, J. G., & Menz, M. H. M. (2017). Consistent behavioural differences between migratory and resident hoverflies. Animal Behaviour, 127, 187–195.

Parker, D. J., Burton, R. R., Diongue-Niang, A., Ellis, R. J., Felton, M., Taylor, C. M., Thorncroft, C. D., Bessemoulin, P., & Tompkins, A. M. (2005). The diurnal cycle of the West African monsoon circulation. Quarterly Journal of the Royal Meteorological Society: A Journal of the Atmospheric Sciences, Applied Meteorology and Physical Oceanography, 131(611), 2839–2860.

Pasanen, M. (2020). Characterization of Pectobacterium strains causing soft rot and blackleg of potato in Finland.

Pedersen, L., Thorup, K., & Tøttrup, A. P. (2019). Annual GPS tracking reveals unexpected wintering area in a long-distance migratory songbird. Journal of Ornithology 160, 1, pp. 265–270. 10.1007/s10336-018-1610-8

Pérez-Bañón, C., Juan, A., Petanidou, T., Marcos-García, M. A., & Crespo, M. B. (2003). The reproductive ecology of Medicago citrina (Font Quer) Greuter (Leguminosae): a bee-pollinated plant in Mediterranean islands where bees are absent. Plant Systematics and Evolution, 241(1), 29–46.

Pérez-Bañón, C., Petanidou, T., & Marcos-García, M. (2007). Pollination in small islands by occasional visitors: the case of Daucus carota subsp. commutatus (Apiaceae) in the Columbretes archipelago, Spain. Plant Ecology, 192(1), 133–151.

Pineda, A., & Marcos-García, M. (2008). Evaluation of several strategies to increase the residence time of Episyrphus balteatus (Diptera, Syrphidae) releases in sweet pepper greenhouses. Annals of Applied Biology, 152(3), 271–276.

Rader, R., Cunningham, S. A., Howlett, B. G., & Inouye, D. W. (2020). Non-bee insects as visitors and pollinators of crops: Biology, ecology, and management. Annual Review of Entomology, 65, 391–407.

Rader, R., Edwards, W., Westcott, D. A., Cunningham, S. A., & Howlett, B. G. (2011). Pollen transport differs among bees and flies in a human-modified landscape. Diversity and Distributions, 17(3), 519–529.

Raftery, A. E., Zimmer, A., Frierson, D. M. W., Startz, R., & Liu, P. (2017). Less than 2 C warming by 2100 unlikely. Nature Climate Change, 7(9), 637–641.

Raymond, L., Sarthou, J.-P., Plantegenest, M., Gauffre, B., Ladet, S., & Vialatte, A. (2014). Immature hoverflies overwinter in cultivated fields and may significantly control aphid populations in autumn. Agriculture, Ecosystems & Environment, 185, 99–105.

Raymond, M., & Pasteur, N. (1996). Evolution of insecticide resistance in the mosquito Culex pipiens: The migration hypothesis of amplified esterase genes. Molecular genetics and evolution of pesticide resistance. 645, pp. 90–96).

Riad, M. H., Scoglio, C. M., McVey, D. S., & Cohnstaedt, L. W. (2017). An individual-level network model for a hypothetical outbreak of Japanese encephalitis in the USA. Stochastic environmental research and risk assessment, 31(2), 353–367. 10.1007/s00477-016-1353-0

Ritchie, S. A., & Rochester, W. (2001). Wind-blown mosquitoes and introduction of Japanese encephalitis into Australia. Emerging Infectious Diseases, 7(5), 900.

Rojo, S., FS, G., Marcos-García, M., Nieto, J. M., & Mier Durante, M. P. (2003). A World Review of Predatory Hoverflies (Diptera, Syrphidae: Syrphinae) and their Prey. CIBIO Ediciones.

Roth, S. K., & Lindroth, R. L. (1994). Effects of CO 2-mediated changes in paper birch and white pine chemistry on gypsy moth performance. Oecologia, 98, 133–138.

Roth, S. K., & Lindroth, R. L. (1995). Elevated atmospheric CO2: effects on phytochemistry, insect performance and insect-parasitoid interactions. Global Change Biology, 1(3), 173–182.

Runge, C. A., Martin, T. G., Possingham, H. P., Willis, S. G., & Fuller, R. A. (2014). Conserving mobile species. Frontiers in Ecology and the Environment, 12(7), 395–402.

Sainz-Borgo, C., Miranda, J., & Lentino, M. (2020). Composition of bird community in Portachuelo Pass (Henri Pittier National Park, Venezuela). Journal of Caribbean Ornithology, 33, 1–14.

Samplonius, J. M., Atkinson, A., Hassall, C., Keogan, K., Thackeray, S. J., Assmann, J. J., Burgess, M. D., Johansson, J., Macphie, K. H., & Pearce-Higgins, J. W. (2021). Strengthening the evidence base for temperature-mediated phenological asynchrony and its impacts. Nature Ecology & Evolution, 5(2), 155–164.

Sánchez-Bayo, F., & Wyckhuys, K. A. G. (2019). Worldwide decline of the entomofauna: A review of its drivers. Biological Conservation, 232, 8–27.

Satterfield, D. A., Sillett, T. S., Chapman, J. W., Altizer, S., & Marra, P. P. (2020). Seasonal insect migrations: massive, influential, and overlooked. Frontiers in Ecology and the Environment, 18(6), 335–344.

Shannon, H. J. (1926). A preliminary report on the seasonal migrations of insects. Journal of the New York Entomological Society, 34(2), 199–205.

Shpedt, A. A., Aksenova, Yu. V., Shayakhmetov, M. R., Zhulanova, V. N., Rassypnov, V. A., & Butyrin, M. V. (2019). Soil and Ecological evaluation of agro-chernozems of Siberia. International Transaction Journal of Engineering, Management, & Applied Sciences & Technologies, 309–318.

Snow, D. W., & Ross, K. F. A. (1952). Insect migration in the Pyrenees. Entomol Mon Mag, 88, 1–6.

Southwood, T. R. E., & Jepson, W. F. (1962). Studies on the populations of Oscinella frit L.(Dipt: Chloropidae) in the oat crop. The Journal of Animal Ecology, 481–495.

Sparks, A. N., Jackson, R. D., Carpenter, J. E., & Muller, R. A. (1986). Insects captured in light traps in the Gulf of Mexico. Annals of the Entomological Society of America, 79(1), 132–139.

Sparks, T. H., Dennis, R. L. H., Croxton, P. J., & Cade, M. (2007). Increased migration of Lepidoptera linked to climate change. European Journal of Entomology, 104(1), 139–143.

Sparks, T. H., Roy, D. B., & Dennis, R. L. H. (2005). The influence of temperature on migration of Lepidoptera into Britain. Global Change Biology, 11(3), 507–514.

Stefanescu, C., Páramo, F., Åkesson, S., Alarcón, M., Ávila, A., Brereton, T., Carnicer, J., Cassar, L. F., Fox, R., Heliölä, J., Hill, J. K., Hirneisen, N., Kjellén, N., Kühn, E., Kuussaari, M., Leskinen, M., Liechti, F., Musche, M., Regan, E. C., … Chapman, J. W. (2013). Multi-generational long-distance migration of insects: Studying the painted lady butterfly in the Western Palaearctic. Ecography 36, 4, pp. 474–486. 10.1111/j.1600-0587.2012.07738.x

Talavera, G., García-Berro, A., Talla, V.N., Ng’iru, I., Bahleman, F., Kébé, K., Nzala, K.M., Plasencia, D., Marafi, M.A., Kassie, A. and Goudégnon, E.O., 2023. The Afrotropical breeding grounds of the Palearctic-African migratory painted lady butterflies (Vanessa cardui). Proceedings of the National Academy of Sciences, 120(16), p.e2218280120.

Tallamy, D. W., & Shriver, W. G. (2021). Are declines in insects and insectivorous birds related? The Condor, 123(1), duaa059.

Tsacas, L. (1984). Nouvelles données sur la biogéographie et l’évolution du groupe Drosophila melanogaster en Afrique. Description de six nouvelles espèces (Diptera, Drosophilidae). Annales de La Société Entomologique de France, 20, 419–438.

Verhelst, B., Jansen, J., & Vansteelant, W. (2011). South West Georgia: an important bottleneck for raptor migration during autumn. Ardea, 99(2), 137–146.

Warren, T. L., Giraldo, Y. M., & Dickinson, M. H. (2019). Celestial navigation in Drosophila. Journal of Experimental Biology, 222(Suppl_1), jeb186148.

Weir, P. T., & Dickinson, M. H. (2012). Flying Drosophila orient to sky polarization. Current Biology, 22(1), 21–27.

Westmacott, H. M., & Williams, C. B. (1954). A migration of Lepidoptera and Diptera in Nepal. Entomologist, 87, 232–234.

Wiegmann, B. M., Trautwein, M. D., Winkler, I. S., Barr, N. B., Kim, J.-W., Lambkin, C., Bertone, M. A., Cassel, B. K., Bayless, K. M., & Heimberg, A. M. (2011). Episodic radiations in the fly tree of life. Proceedings of the National Academy of Sciences, 108(14), 5690–5695.

Wikelski, M., Moskowitz, D., Adelman, J. S., Cochran, J., Wilcove, D. S., & May, M. L. (2006). Simple rules guide dragonfly migration. Biology Letters, 2(3), 325–329.

Williams, C. B. (1958). Insect Migration. New Naturalist 36. Collins.

Williams, C. B., Common, I. F. B., French, R. A., Muspratt, V., & Williams, M. C. (1956). Observations on the migration of insects in the Pyrenees in the autumn of 1953. Transactions of the Royal Entomological Society of London, 108(9), 385–407.

Wolf, W.W., Sparks, A.N., Pair, S.D., Westbrook, J.K. and Truesdale, F.M., 1986. Radar observations and collections of insects in the Gulf of Mexico. Insect flight: dispersal and migration (pp. 221–234). Springer Berlin Heidelberg.

Wotton, K. R., Gao, B., Menz, M. H. M., Morris, R. K. A., Ball, S. G., Lim, K. S., Reynolds, D. R., Hu, G., & Chapman, J. W. (2019). Mass seasonal migrations of hoverflies provide extensive pollination and crop protection services. Current Biology, 29(13), 2167–2173.

Zeng, J., Liu, Y., Zhang, H., Liu, J., Jiang, Y., Wyckhuys, K. A. G., & Wu, K. (2020). Global warming modifies long-distance migration of an agricultural insect pest. Journal of Pest Science, 93, 569–581.

Караџић, В. С. (2005). Живот и обичаји народа српскога (Issue 9). Политика

